# PVN-mPFC OT projections modulates pup-directed pup care or attacking in virgin mandarin voles

**DOI:** 10.1101/2024.03.06.583718

**Authors:** Lu Li, Yin Li, Caihong Huang, Wenjuan Hou, Zijian Lv, Lizi Zhang, Yishan Qu, Yahan Sun, Kaizhe Huang, Xiao Han, Zhixiong He, Fadao Tai

## Abstract

In many species, adult animals may exhibit caregiving or aggression towards conspecific offspring. The neural mechanisms underlying the infanticide and pup care remain poorly understood. Here, using monogamous virgin mandarin voles (*Microtus mandarinus*) that may exhibit pup care or infanticide, we found that more oxytocin (OT) neurons in the paraventricular nucleus (PVN) were activated during pup caring than infanticide. Optogenetic activation of OT neurons in the PVN facilitated pup-caring in male and female mandarin voles. In infanticide voles, optogenetic activation of PVN OT cells prolonged latency to approach and attack pups, whereas inhibition of these OT neurons facilitated approach and infanticide. In addition, OT release in the medial prefrontal cortex (mPFC) in pup-care voles increased upon approaching and retrieving pups, and decreased in infanticide voles upon attacking pups. Optogenetic activation of PVN OT neuron projections to the mPFC shortened the latency to approach and retrieve pups and facilitated the initiation of pup care, whereas inhibition of these projections had little effect. For pup-care females, neither activation nor inhibition of the fibers affected their behavior towards pups. In infanticide male and female voles, optogenetic activation of PVN-mPFC OT projection fibers prolonged the latency to approach and attack pups and suppressed the initiation of infanticide, whereas inhibition of these projections promoted approach and infanticide. Finally, we found that intraperitoneal injection of OT promoted pup care and inhibited infanticide behavior. It is suggested that the OT system, especially PVN OT neurons projecting to mPFC, modulates pup-directed behaviors and OT can be used to treat abnormal behavioral responses associated with some psychological diseases such as depression and psychosis.

## Introduction

Both paternal and maternal care are critical to the survival as well as the physical and mental well-being of the offspring (He et al., 2019). Although we know a great deal about the neural mechanisms underlying maternal care, the neural substrates of paternal behavior remain elusive because of lacking ideal animal model of paternal care. Only in some monogamous rodents (e.g. prairie voles), canids and primates, do males assist and spend a great deal of energy caring for pups (Malcolm, 2015; Mendoza & Mason, 1986; Rosenfeld, Johnson, Ellersieck, & Roberts, 2013). However, some male rodents without reproductive experience also show paternal care toward alien pups, while some others ignore or even attack alien pups (Dai et al., 2022). These different pup-directed behavioral responses are based on their physiological and environmental states, and the killing of young of the conspecifics by sexually inexperienced mammals is a widespread phenomenon among different animal taxa (Lukas & Huchard, 2014). Infanticide is thought to benefit the infanticide by promoting their own reproduction (Hrdy, 1974). In laboratory mice, male mice without pairing experience typically display infanticide (Svare & Mann, 1981), but males are able to shift from infanticide to pup care when they have the opportunity to encounter their own offspring (Elwood, 1977). Compared to our extensive understanding of the maternal circuit, little is known about the neural substrates underlying female infanticide. The virgin mandarin voles (*Microtus mandarinus*) naturally exhibit biparental care and infanticide in both female and male that provide ideal animal model to reveal mechanism underlying paternal care in male and infanticide in females.

OT is well-known as a key hormone for initiating and maintaining maternal care (Yoshihara, Numan, & Kuroda, 2018), which is primarily synthesized in the PVN and supraoptic nucleus (SON), among which the PVN plays an important role in initiating maternal care in rats (Munetomo, Ishii, Miyamoto, Sakuma, & Kondo, 2016). There is evidence to suggest that OT not only regulates maternal motivation, but also mediates paternal behavior (Bales, Kim, Lewis-Reese, & Sue Carter, 2004). When mouse fathers were exposed to their pups, OT neurons in the PVN were specifically activated and they also showed more aggression towards intruders to protect the own pups (Shabalova et al., 2020). Compared with rodents, similar neuropeptides and hormones are involved in paternal behavior in non-human primates (Woller et al., 2012). Similar to other mammals, paternal care only exists in a few primate species (Dulac, O’Connell, & Wu, 2014; Woller et al., 2012). A study on marmoset monkeys showed that fathers have higher levels of OT secretion in the hypothalamus than non-fathers (Woller et al., 2012), and intraventricular infusion of OT reduces the tendency of marmoset fathers to refuse to transfer food to their young offspring (Saito & Nakamura, 2011). However, where and how pathways of OT neurons regulate pup care and infanticide behavior remain largely unknown.

The mPFC is involved in attention switching, decision-making, behavioral flexibility and planning, making it potentially crucial for rapidly expressing pup care or infanticide behavior. A study on virgin male and female mice found that the mPFC lesion (targeting the prelimbic cortex) significantly affected number of females and males showing pup cares and infanticide. 50% females in lesioned group exhibited maternal while 100% of sham operated groups show maternal care, whereas the 100% lesioned males exhibited infanticide, 83% of control males showed infanticide (Alsina-Llanes & Olazábal, 2021). It has been reported that the mPFC is highly activated in human mothers when they hear cry from their babies (Lorberbaum et al., 2002). The mPFC is also activated when rat mother contacts their pups first time (Fleming & Korsmit, 1996), while the damage of the mPFC disrupts pup retrieving and grooming behavior in rats (Afonso, Sison, Lovic, & Fleming, 2007). In rodents and humans, the mPFC was activated during the process of caring offspring (Hernández-González, Navarro-Meza, Prieto-Beracoechea, & Guevara, 2005; Seifritz et al., 2003). A study in rat mothers indicated that inactivation or inhibition of neurons in the mPFC largely reduced pup retrieval and grouping (Febo, Felix-Ortiz, & Johnson, 2010). In a subsequent study on firing patterns in the mPFC of rat mother suggested that sensory-motor processing carried out in the mPFC may affect decision making of maternal care to their pups (Febo, 2012). Examining different regions of the mPFC (anterior cingulate (Cg1), prelimbic (PrL), infralimbic (IL)) of new mother identified a role for the IL cortex in biased preference decision-making in favour of the offspring (Pereira & Morrell, 2020). A study on rats suggests that the IL and Cg1 subregion in mPFC are the motivating circuits for pup-specific biases in the early postpartum period (Pereira & Morrell, 2011), while the PrL subregion, are recruited and contribute to the expression of maternal behaviors in the late postpartum period (Pereira & Morrell, 2011). In addition, a large number of neurons in the mPFC express oxytocin receptors (OTRs) (Smeltzer, Curtis, Aragona, & Wang, 2006). OT in the circulation of mice can act on the medial prefrontal cortex (mPFC) to increase social interaction and maternal behavior (Munesue et al., 2021). Although there are some studies on the role of the mPFC in pup care, the involvement of mPFC OT projections in pup cares and infanticide requires further research. Thus, we hypothesized that PVN OT neurons projections to mPFC may causally control paternal or infanticide behaviors.

Based on the potential antagonistic effects between pup care and infanticide behavior in neural mechanisms (Dulac et al., 2014; Kohl, Autry, & Dulac, 2017; Mei, Yan, Yin, Sullivan, & Lin, 2023), as well as the potential role of PVN-to-mPFC OT projections in pup care and infanticide. Here, a combination of methods, including immunohistochemistry, optogenetics, fiber photometry and intraperitoneal injection of OT were used to reveal neural mechanisms underlying paternal care and infanticide. We found that PVN OT neurons regulated the expression of pup care and infanticide, and further identified the involvement of PVN-to-mPFC OT projections in paternal care and infanticide. Collectively, these findings establish a regulatory role for PVN-to-mPFC OT neurons in the expression of pup-directed behaviors and suggest potential targets for the future development of intervention strategies against psychiatric disorders associated with infanticide such as depression and psychosis.

## Results

### Pup care behavior activate more OT^+^ cells than infanticide in PVN

In order to observe the activated OT neurons in virgin voles during pup care and infanticide behaviors, we co-stained OT and c-Fos on brain slices from voles exhibiting different behaviors using immunofluorescence method (Fig. 1a). Histological analysis showed no difference in the number of OT or c-Fos positive cells between the pup care and infanticide groups of female (Fig. 1b,c, Fig. 1-source data 1) and male (Fig. 1e,f, Fig. 1-source data 1) voles. Approximately 11% (male) and 14% (female) of OT cells expressed c-Fos during pup caring, whereas only about 3% (males) and 7% (females) of OT neurons were labeled by c-Fos during infanticide (female: t (6) = 5.173, *P* < 0.01, d = 3.658, Fig. 1d, Fig. 1-source data 1; male: t (6) =2.607, *P* < 0.05, d = 1.907, Fig. 1g, Fig. 1-source data 1). In male and female voles, more OT neurons were activated during pup-caring than infanticide (Fig. 1d, g). In addition, females displaying pup care and infanticide showed higher merge rates of OT and c-Fos than males displaying the same behaviors (F(1,12) = 5.002, P = 0.045, η^2^ = 0.294, Fig. 1d, g, Fig. 1-source data 1).

**Figure 1:**
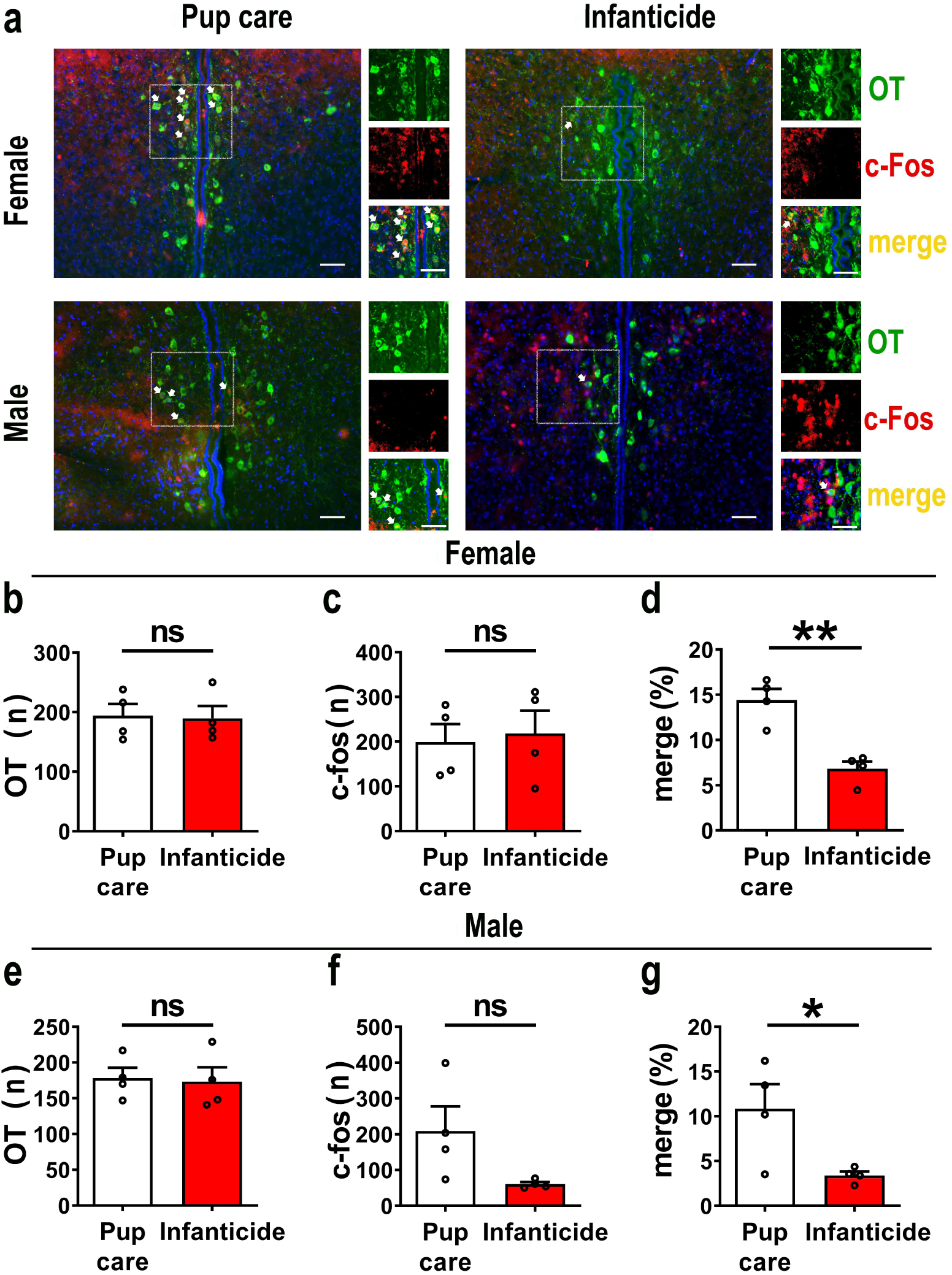
Activated OT neurons in the PVN of mandarin voles during pup care (n =4) and infanticide (n = 4). a. Representative histological images of OT (green) and c-Fos (red) positive cells in PVN: Blue, DAPI: Yellow, merged cells. Enlarged views of the boxed area were shown to the right of each image, white arrows indicated the overlap of OT and c-fos positive cells. Objective: 20x. Scale bars, 50 μm. b. Number of OT positive cells in pup care and infanticide female voles. c. Number of c-Fos positive cells in pup care and infanticide female voles. d. Percentage of c-Fos-expressing cells in OT cells of PVN from pup care and infanticide female voles. \*\**P* < 0.01. Independent sample t-tests. e. Number of OT cells in pup care and infanticide male voles. f. Number of c-Fos cells in pup care and infanticide male voles. g. Percentage of c-Fos-expressing cells in OT cells of PVN from pup care and infanticide male voles. \**P* < 0.05. Independent sample t-tests. Erroe bars indicate SEM. Figure 1-souce data 1. Statistical results of the number of OT-positive cells, the number of c-Fos-positive cells, and the percentage of OT and c-Fos merged neurons in the total PVN OT neurons in female and male pup-care and infanticide voles.

### Effects of optogenetic activation of PVN OT neurons on pup-directed behaviors

To reveal causal role of PVN OT neuron in regulation of pup-care and infanticide behaviors, effects of optogenetic activation of PVN OT neurons on pup-directed behaviors were investigated (Fig. 2 a,b,c). Over 89% of CHR2 expression overlapped with OT neurons indicating high specificity of CHR2 virus (Fig. 2d, Figure 2-source data 1). 473 nm light stimulation increased c-Fos expression in CHR2 virus infected brain region that validated the effect of optogenetic activation (Supplementary Data Fig.1a-c). We found that optogenetic activation of PVN OT cells significantly reduced latency to approach (CHR2: off vs on F (1, 7) = 11.374, *P* < 0.05, OFF/ON: η^2^ = 0.592, Fig. 2o, Figure 2-source data 1) and retrieve pups (CHR2: off vs on F (1, 4) = 14.755, *P* < 0.05, OFF/ON: η^2^ = 0.156, Fig. 2p, Figure 2-source data 1) and prolonged crouching time (CHR2: off vs on F (1, 7) = 60.585, *P* < 0.001, OFF/ON: η^2^ = 0.419, Fig. 2s, Figure 2-source data 1) in pup-care males, but had no effect on females (Fig. 2j-n, Figure 2-source data 1), nor in control virus group. Optogenetic activation of these neurons significantly reduced the latency to approach and attack pups in male (approach: CHR2: off vs on F (1, 5) = 185.509, *P* < 0.0001, OFF/ON: η^2^ = 0.552, Fig. 2h, Figure 2-source data 1; infanticide: CHR2: off vs on F (1, 5) = 59.877, *P* < 0.01, OFF/ON: η^2^ = 0.526, Fig. 2i, Figure 2-source data 1) and female voles (approach: CHR2: off vs on F (1, 6) = 64.810, *P* < 0.001, OFF/ON: η^2^ = 0.915, Fig. 2f, Figure source data 1; infanticide: CHR2: off vs on F (1, 6) = 75.729, *P* < 0.001, OFF/ON: η^2^ = 0.940, Fig. 2g, Figure 2-source data 1) displaying infanticide behaviors whereas had no effect on control virus group (Fig. 2f-I, Figure 2-source data 1). Further, we conducted a two-way rmANOVA on the CHR2 data set for both sexes and found that pup-care females exhibited shorter latencies to approach (OFF: gender simple effect F(1,13) = 5.735, P = 0.032, η^2^ = 0.306, Fig. 2j, o) and retrieve pups (OFF: gender simple effect F(1,10) = 13.040, P = 0.005, η^2^ = 0.566, Fig. 2k, p) than males (Figure 2-source data 1). These results suggest that activation of PVN OT neurons facilitate pup care behavior and significantly inhibit infanticide behavior.

**Fig. 2.**
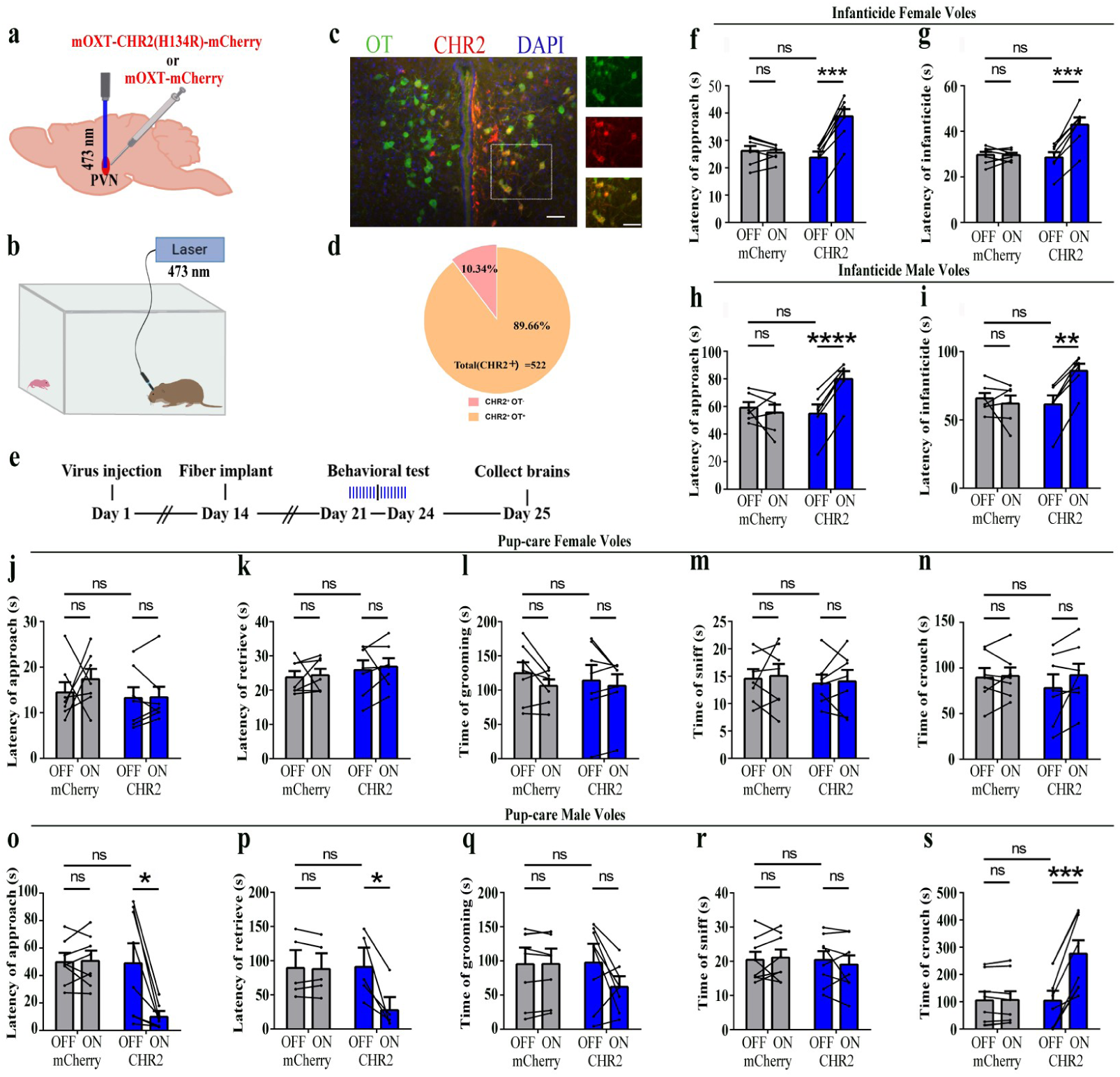
Effects of optogenetic activation of PVN OT neurons on pup-directed behaviors. a. Schematic of virus injection and optical fiber implantation. b. Schematic diagram of the behavioral test. c. Representative histologic images of CHR2 (red) expression and OT staining (green), enlarged view of the boxed area is on the right side. Blue, DAPI. Objective: 20x. Scale bars, 50 μm. d. Statistics on the specificity of CHR2 expression in 3 voles, more than 89% of CHR2 positive neurons overlapped with OT positive neurons. e. Time line of the experiment. f-i, Approach (f, h) and infanticide (g, i) latency of infanticide 7 female and 6 male voles in mCherry (control virus group) and CHR2 groups. \*\**P* < 0.01 vs. CHR2 OFF; \*\*\**P* < 0.001 vs. CHR2 OFF; \*\*\*\**P* < 0.0001 vs. CHR2 OFF. Two-way rmANOVA (factors: treatment × stimulus). j-n, Latency to approach (j), latency to retrieve (k), grooming time (l), sniffing time (m) and crouching time (n) of 7 pup-care female voles in female control virus and CHR2 groups. o-s, Latency to approach (o), latency to retrieve (p), grooming time (q), sniffing time (r) and crouching time (s) of 8 pup-care male voles in male control virus and CHR2 groups. \**P* < 0.05 vs. CHR2 OFF, \*\*\**P* < 0.001 vs. CHR2 OFF. Two-way rmANOVA (factors: treatment × stimulus). Error bars indicate SEM. Figure 2-source data 1. Statistical results of the number of cells expressing only CHR2 and co-expressing CHR2 and OT in the PVN of 3 voles with injection of optogenetic virus, the latency to approach and attack pups in infanticide voles, and the latency to approach, retrieve, duration of grooming, sniffing, and crouching in pup-care voles.

### Effects of optogenetic inhibition of PVN OT neurons on pup-directed behaviors

To further verify the roles of PVN OT neurons on pup-induced behavior, we optogenetically inhibited OT cells by eNpHR virus and tested pup-directed behaviors (Fig. 3a,b,c). More than 90% of neurons expressing eNpHR overlapped with OT positive neurons indicating high specificity of eNpHR virus infection (Fig. 3d, Figure 3-source data 1). 589 nm light stimulation to eNpHR virus infected brain regions reduced c-Fos expression verifying the effectiveness of opotogenetic inhibition via eNpHR virus (Supplementary Data Fig. 1d-f). Inhibition of PVN OT neurons showed no significant effect on pup care behavior in male and female voles that spontaneously exhibited pup caregiving behaviors (Fig. 3j-s, Figure 3-source data 1). For both male and female voles in the infanticide group, optogenetic inhibition significantly shortened the latency to approach (female: F (1, 5) = 1331.434, *P* < 0.0001, OFF/ON: η^2^ = 0.980, Fig. 3f, Figure 3-source data 1; male: F (1, 5) = 10.472, *P* < 0.05, OFF/ON: η^2^ = 0.690, Fig. 3h, Figure 3-source data 1) and attack pups (female: F (1, 5) = 291.606, *P* < 0.0001, OFF/ON: η^2^ = 0.991, Fig. 3g, Figure source data 1; male: F (1, 5) = 46.901, *P* < 0.01, OFF/ON: η^2^ = 0.837, Fig. 3i, Figure 3-source data 1). In addition, we performed a two-way rmANOVA on the eNpHR group data for both sexes and found that pup-care females exhibited shorter latency to approach (gender main effect F(1,10) = 62.131, P < 0.0001, η^2^ = 0.861, Fig. 3j, o) and retrieve (gender main effect F(1,10) = 137.393, P < 0.0001, η^2^ = 0.932, Fig, 3k, p) than males (Figure 3-source data 1). These results suggest that inhibition of OT neurons in the PVN significantly facilitates infanticide behavior.

**Fig. 3.**
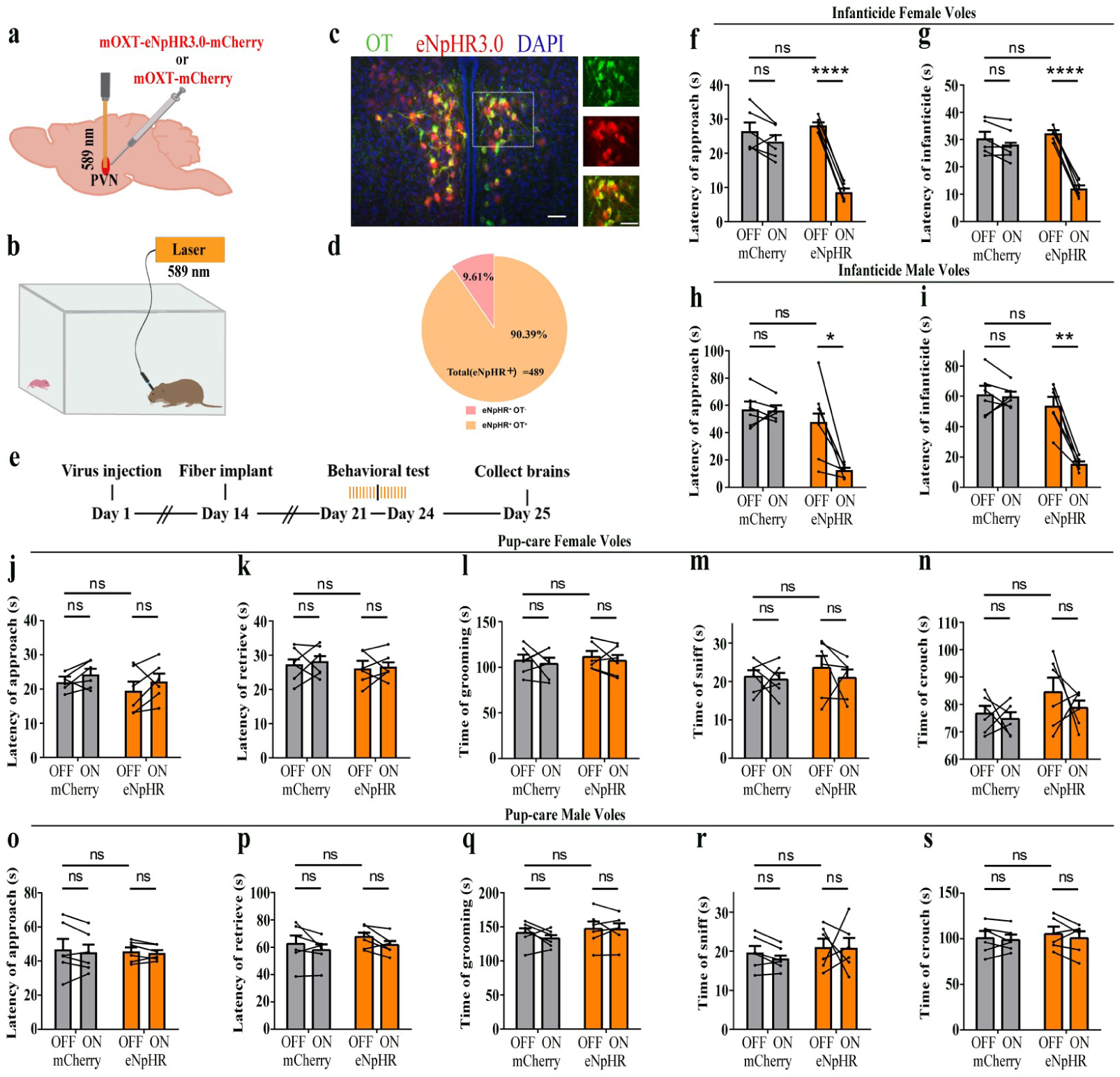
Effects of optogenetic inhibition of PVN OT neurons on pup-directed behaviors. a. Schematic of virus injection and optical fiber implantation. b. Schematic diagram of the behavior test. c. Representative histological images of OT staining (green) and eNpHR (red) expression, enlarged view of the boxed area is shown on the right side. Blue, DAPI. Objective: 20x. Scale bars, 50 μm. d. Statistics on the specificity of CHR2 expression in 3 voles, more than 90% of eNpHR expression overlapped with OT. e. Time line of the experiment. f-i, Approach (f, h) and infanticide (g, i) latency in 6 female (f, g) and 6 male (h, i) infanticide voles. \**P* < 0.05 vs. eNpHR OFF; \*\**P* < 0.01 vs. eNpHR OFF; \*\*\*\**P* < 0.0001 vs. eNpHR OFF. Two-way rmANOVA (factors: treatment × stimulus). j-n, Approach latency (j), retrieval latency (k), grooming time (l), sniffing time (m) and crouching time (n) in 6 control mCherry and eNpHR group of pup-care female voles. o-s, Approach latency (o), retrieval latency (p), grooming time (q), sniffing time (r) and crouching time (s) in 6 control mCherry and eNpHR group of pup-care male voles. Error bars indicate SEM. Figure 3-source data 1. Statistical results of the number of cells expressing only eNpHR and co-expressing eNpHR and OT in the PVN of 3 voles with injection of optogenetic virus, the latency to approach and attack pups in infanticide voles, and the latency to approach, retrieve pups, duration of grooming, sniffing, and crouching in pup-care voles.

### Changes in OT release upon pup-directed behaviors

The results of the optogenetic manipulation demonstrated that PVN OT neurons regulated pup-induced behavior. We next detected the OT release in the mPFC during pup-induced behavior by OT1.0 sensor (Fig. 4a-c). Pup-caring female and male voles showed a significant increase (female: F (1.958, 13.708) = 45.042, *P* < 0.001, η^2^ = 0.865; male: F (5, 35) = 24.057, *P* < 0.01, η^2^ = 0.775) in the signal for OT1.0 sensors upon approaching (female: *P* < 0.01; male: *P* < 0.05) and retrieving (female: *P* < 0.01; male: *P* < 0.05), whereas there was no significant difference in the signal at the onset of other behaviors compared with the signal before the introduction of the pups (Fig. 4f-m, Figure 4-source data 1). In addition, we compared the signals of OT1.0 sensors in the pup-caring voles upon the first, second and third approaches to the pups (female: F (2, 14) = 10.917, *P* < 0.01, η^2^ = 0.609; male: F (2, 14) = 13.351, *P* < 0.01, η^2^ = 0.656), OT release peaked at the first approach and tended to decrease thereafter (Fig. 4d,e, Figure 4-source data 1). In infanticide female and male voles, OT release decreased upon attacking in infanticide males (F (1.117, 7.822) = 85.803, *P* < 0.001, η^2^ = 0.838) and females (F (1.068, 7.479) =36.336, *P* < 0.001, η^2^ = 0.925) (Fig. 4n-r, Figure 4-source data 1). In addition, no significant changes in signals were detected from the individuals with control AAV2/9-hSyn-OTmut during pup-directed behaviors (Supplementary Data Fig. 2). In addition, no changes in OT release were detected while subjects were exposed to object with similar size and shape as the pup (Supplementary Data Fig. 3). Besides, we found that pup-care females showed higher AUC per second than males during approaching (gender simple effect F(1,14) = 27.740, P = 0.000119, η^2^ = 0.665, Fig. 4l, m, Figure 4-source data 1) and retrieving (gender simple effect F(1,14) = 11.695, P = 0.004, η^2^ = 0.455, Fig. 4l, m, Figure 4-source data 1) the pups. These results indicate that mPFC OT release significantly increased upon approaching and retrieving in pup-care voles, but decreased upon attacking pups in infanticide voles.

**Fig. 4.**
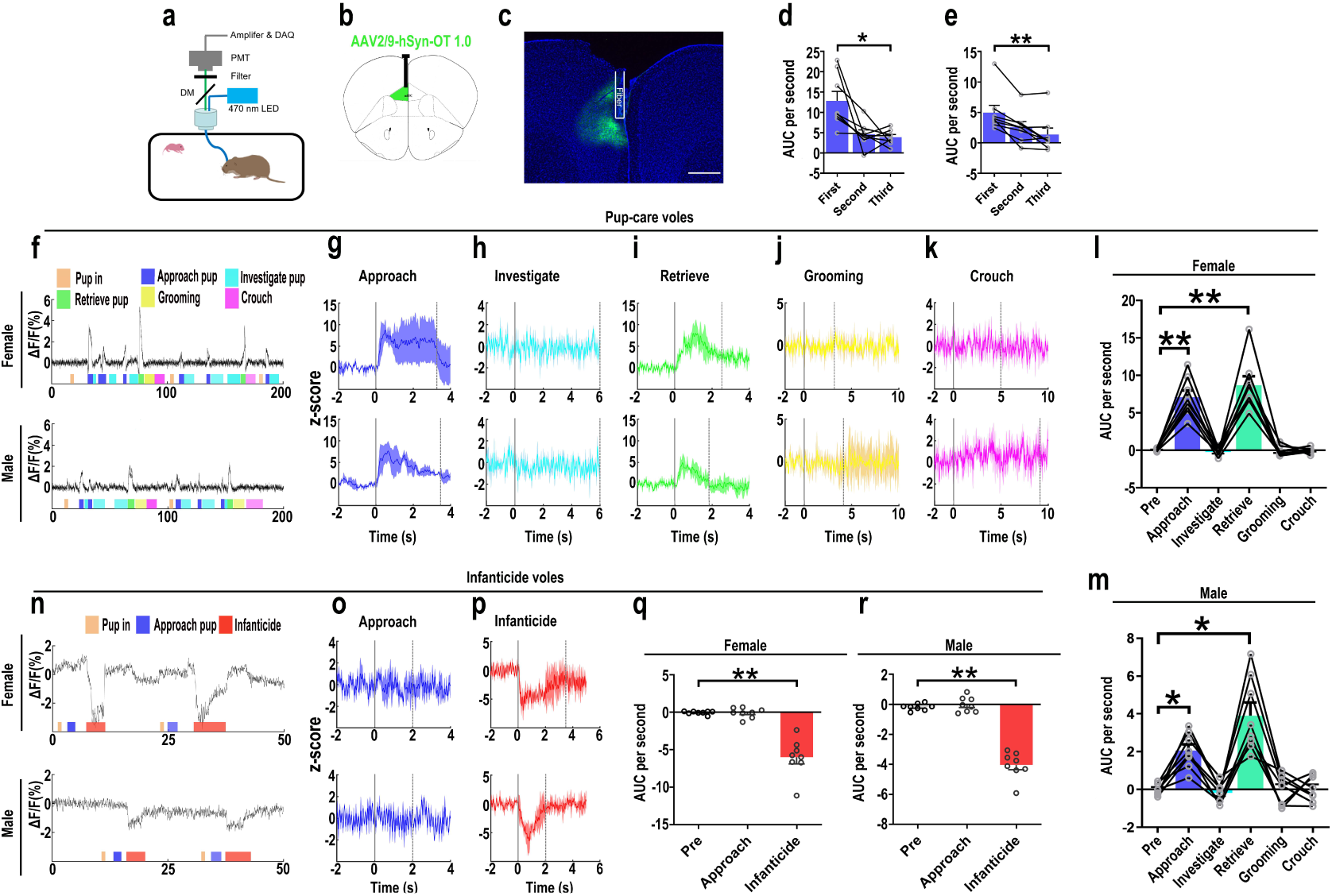
OT release in the mPFC upon pup-directed behaviors. a. Recording instrument settings. b. Illustrations of viral expression and optical fibre location. c. Representative histological image of OT1.0 sensor (green) and optical fibre locations. Blue, DAPI. Objective: 4x. Scale bars, 500 μm. d, e, Area under the curve (AUC) per second for pup-care female (d) and male (e) voles approaching pups for the first, second and third time (n = 8). \**P* < 0.05 vs. first. \*\**P* < 0.01 vs. first. One-way rmANOVA. f. Representative ΔF/F traces in pup-care female (f, top) and male (f, bottom) voles during interaction with pups. g-k, Post-event histograms (PETHs) of z-score of OT1.0 sensor for the following pup-directed behaviors: approach (g), investigate (h), retrieve (i), grooming (j) and crouch (k). l,m, The mean AUC of z-scores for pup-care female (l) and male (m) voles across various pup-directed behaviors (n = 8). Female: \*\**P* < 0.01 vs approach. *P* < 0.01 vs retrieve. Male: \**P* < 0.05 vs approach. *P* < 0.05 vs retrieve. One-way rmANOVA. n. Representative ΔF/F traces in infanticide female (n, top) and male (n, bottom) voles during interaction with pups. o,p, PETHs of z-score of OT1.0 sensor for approach and infanticide in infanticide voles. q,r, The mean AUC of z-score of OT1.0 sensor for pre-pup exposure, approach and infanticide in infanticide female (q) and male (r) voles (n = 8). \*\**P* < 0.01 vs infanticide. One-way rmANOVA. Eerro bars indicate SEM. Figure 4-source data 1. Area under the curve per second for pre-pup exposure, approach, and infanticide behaviors in infanticide voles and area under the curve per second for first, second, and third approaches to pups, as well as pre-pup exposure, approach, investigation, retrieval, grooming, and crouching behaviors in pup-care voles.

### Effects of optogenetic activation of PVN OT neurons fibers in the mPFC on pup-directed behaviors

Previous experiment found that OT release in the mPFC changed upon pup-directed behavior. To manipulate the neural circuit, we first verified oxytocin projections from PVN to the mPFC. We injected retrogradely labeled virus in the mPFC and observed the overlap of virus with OT in the PVN (Fig. 5), and we also counted the PVN OT neurons projecting to mPFC and found that approximately 45.16% and 40.79% of cells projecting from PVN to the mPFC were OT-positive, and approximately 18.48% and 18.89% of OT cells in the PVN projected to the mPFC in females and males, respectively (Supplementary Data Fig. 4). Then, we tested whether optogenetic activation of the PVN OT neuron projection fibers in the mPFC affects pup-induced behaviors (Fig. 6a-d). Similar to previous results, in the pup caring group, activation of the fibers facilitated approaching (F (1,5) = 23.915, *P* < 0.01, OFF/ON: η^2^ = 0.760) and retrieving (F (1,5) = 39.664, *P* < 0.01, OFF/ON: η^2^ = 0.907) in male voles (Fig. 6o-s, Figure 6-source data 1), but had no effect on females (Fig. 6j-n, Figure 6-source data 1). In male and female infanticide group voles, optogenetic activation of the PVN OT neuron projection fibers prolonged latency to approach (female: F (1,5) = 37.094, *P* < 0.01, OFF/ON: η^2^ = 0.875; male: F (1,5) = 74.718, *P* < 0.001, OFF/ON: η^2^ = 0.889) and attack pups (female: F (1,5) = 38.347, *P* < 0.01, OFF/ON: η^2^ = 0.877; male: F (1,5) = 61.589, *P* < 0.01, OFF/ON: η^2^ = 0.910) (Fig. 6f-i, Figure 6-source data 1). These results suggest that activation of the PVN OT neurons to mPFC projection promoted the onset of pup care behavior in pup-care male voles and inhibited the onset of infanticide behavior in infanticide voles.

**Fig. 5.**
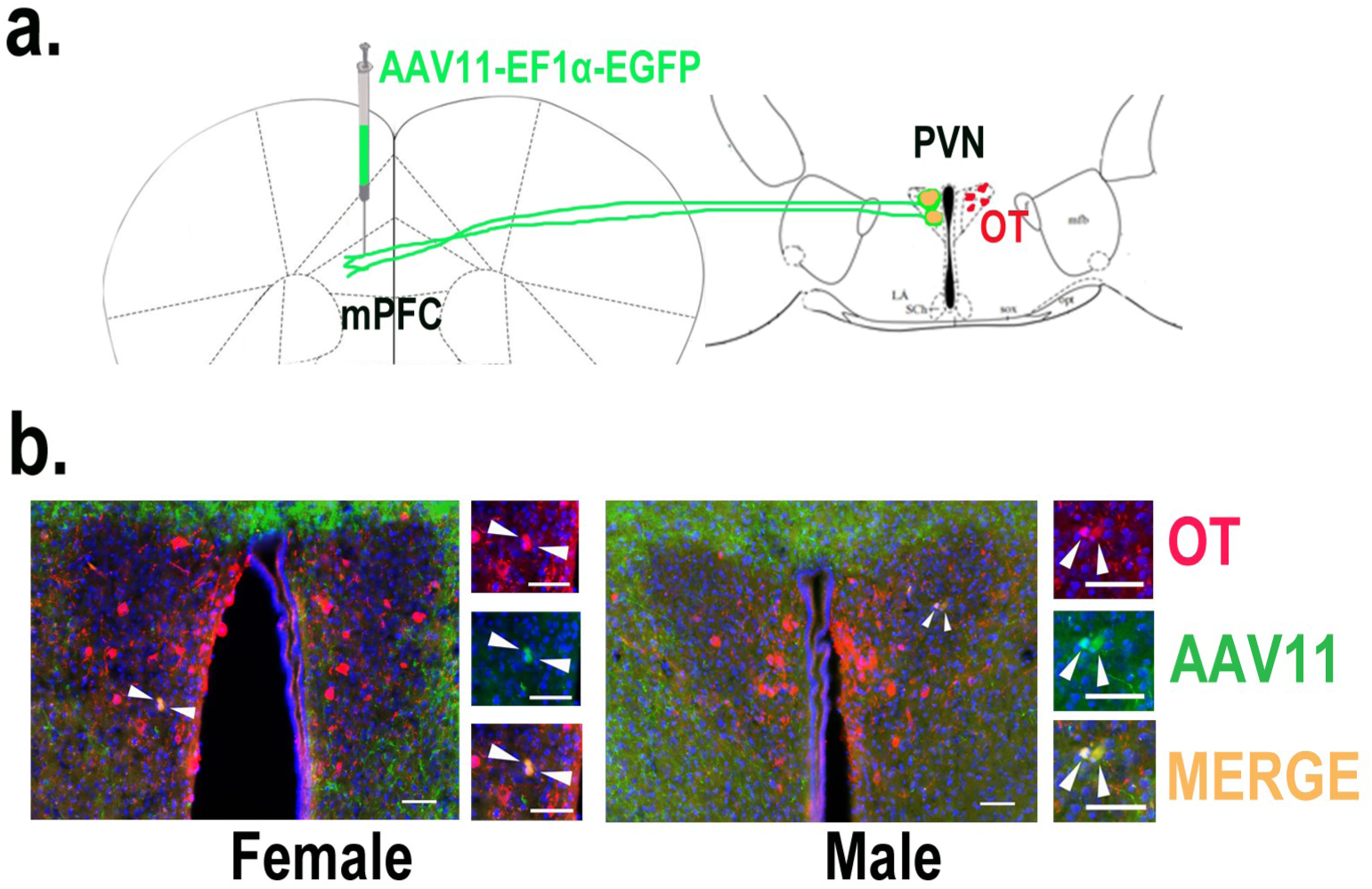
Determination of PVN to mPFC oxytocin projection. a. Schematic diagram of mPFC virus injection and OT staining. b. Histological pictures of OT (red) and AAV11 (green) co-staining in male and female. Yellow, merged. Blue, DAPI. Objective: 20x. Scale bars, 50 μm.

**Fig. 6.**
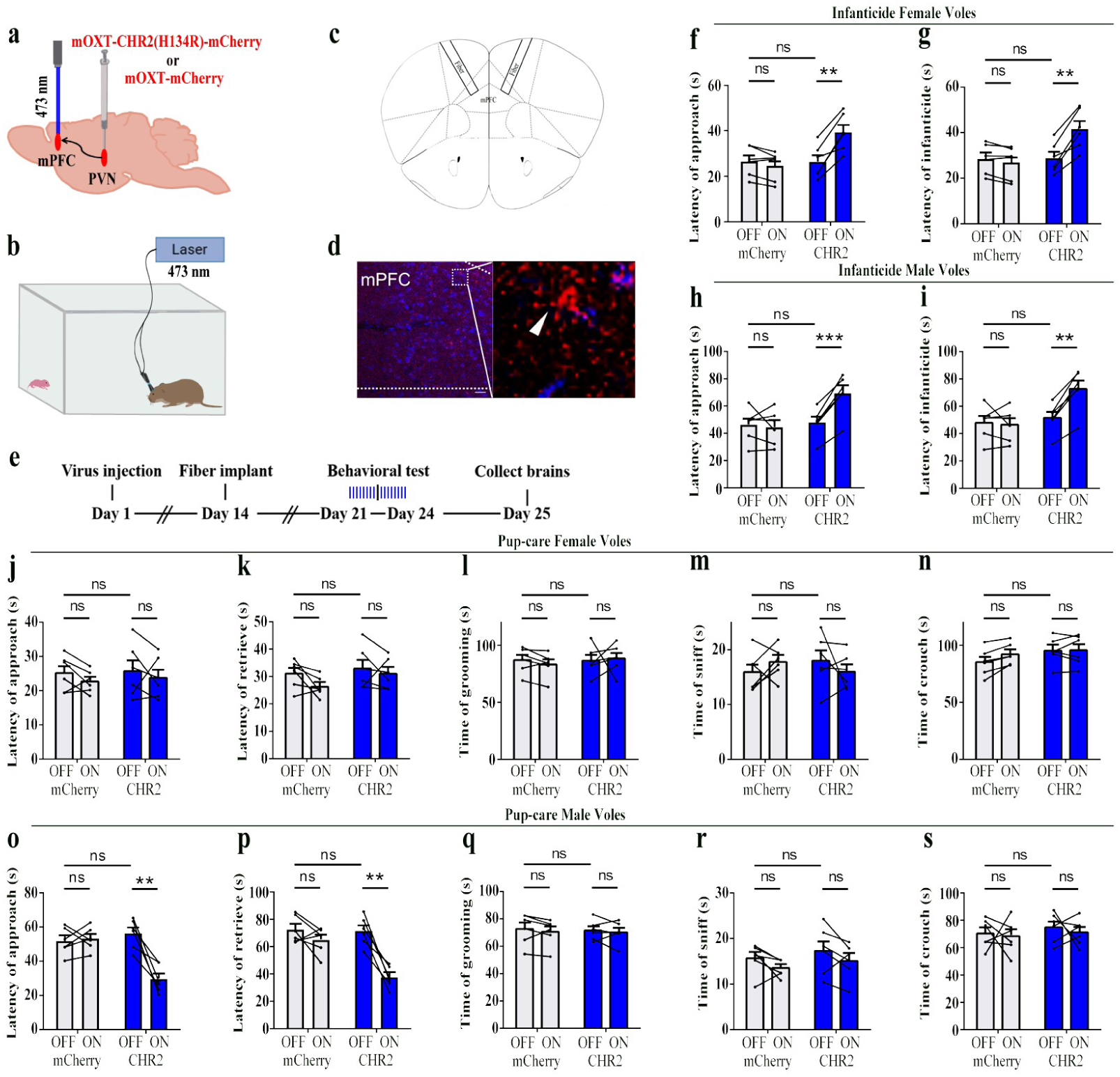
Effects of optogenetic activation of the PVN OT neuron projection fibers on pup-directed behaviors. a. Schematic of virus injection and optical fiber implantation. b. Schematic diagram of the behavior test. c. Illustration of optical fiber implantation in the target brain region. d. Representative histological pictures of the fiber position and projection fibers. Blue, DAPI. Objective: 20x. Scale bars, 50 μm. e. Time line of the experiment. f-i, Changes in approach latency (f, h) and infanticide latency (g, i) of female (f, g) and male (h, i) infanticide voles in CHR2 and control mCherry group before and after delivery of light (n = 6). \*\**P* < 0.01 vs. CHR2 OFF; \*\*\**P* < 0.001 vs. CHR2 OFF. Two-way ANOVA (factors: treatment × stimulus). j-n, Approach latency (j), retrieval latency (k), grooming time (l), sniffing time (m) and crouching time (n) in control mCherry and CHR2 group of pup-care female voles (n = 6). o-s, Approach latency (o), retrieval latency (p), grooming time (q), sniffing time (r) and crouching time (s) in control mCherry and CHR2 group of pup-care male voles (n = 6). \*\**P* < 0.01 vs. CHR2 OFF. Two-way rmANOVA (factors: treatment × stimulus). Error bars indicate SEM. Figure 6-source data 1. Statistical results of the latency to approach and attack pups in infanticide voles, and the latency to approach, retrieve pups, duration of grooming, sniffing, and crouching in pup-care voles.

### Optogenetic inhibition of the PVN OT neuron projection fibers promoted infanticide

We then optogenetically suppressed the projection fibers from PVN OT neurons to mPFC and observed changes in pup-directed behaviors (Fig. 7a-d). Similar to the results of the PVN OT neurons inhibition, we found that optogenetic inhibition of the PVN OT neuron projection fibers promoted approach (F (1,5) = 119.093, *P* < 0.001, OFF/ON: η^2^ = 0.877) and infanticide (F (1,5) = 112.501, *P* < 0.001, OFF/ON: η^2^ = 0.885) in infanticide male (Fig. 7h,i, Figure 7-source data 1) and female voles (approach: F (1,5) = 280.031, *P* < 0.0001, OFF/ON: η^2^ = 0.853; infanticide: F (1,5) = 268.694, *P* < 0.0001, OFF/ON: η^2^ = 0.838) (Fig. 7f,g, Figure 7-source data 1). For pup-care male and female voles, inhibition of these fibers did not significantly affect their pup care behaviors (Fig. 7j-s, Figure 7-source data 1). To validate the effectivity of fiber optogenetic inhibition, we combined optogenetic inhibition with OT1.0 sensors and recorded a decrease in OT release upon inhibition of fibers (Supplementary Data Fig. 5). These results suggest that optogenetic inhibition of the PVN OT neuron projection fibers promotes the onset of infanticide behavior in infanticide voles.

**Fig. 7.**
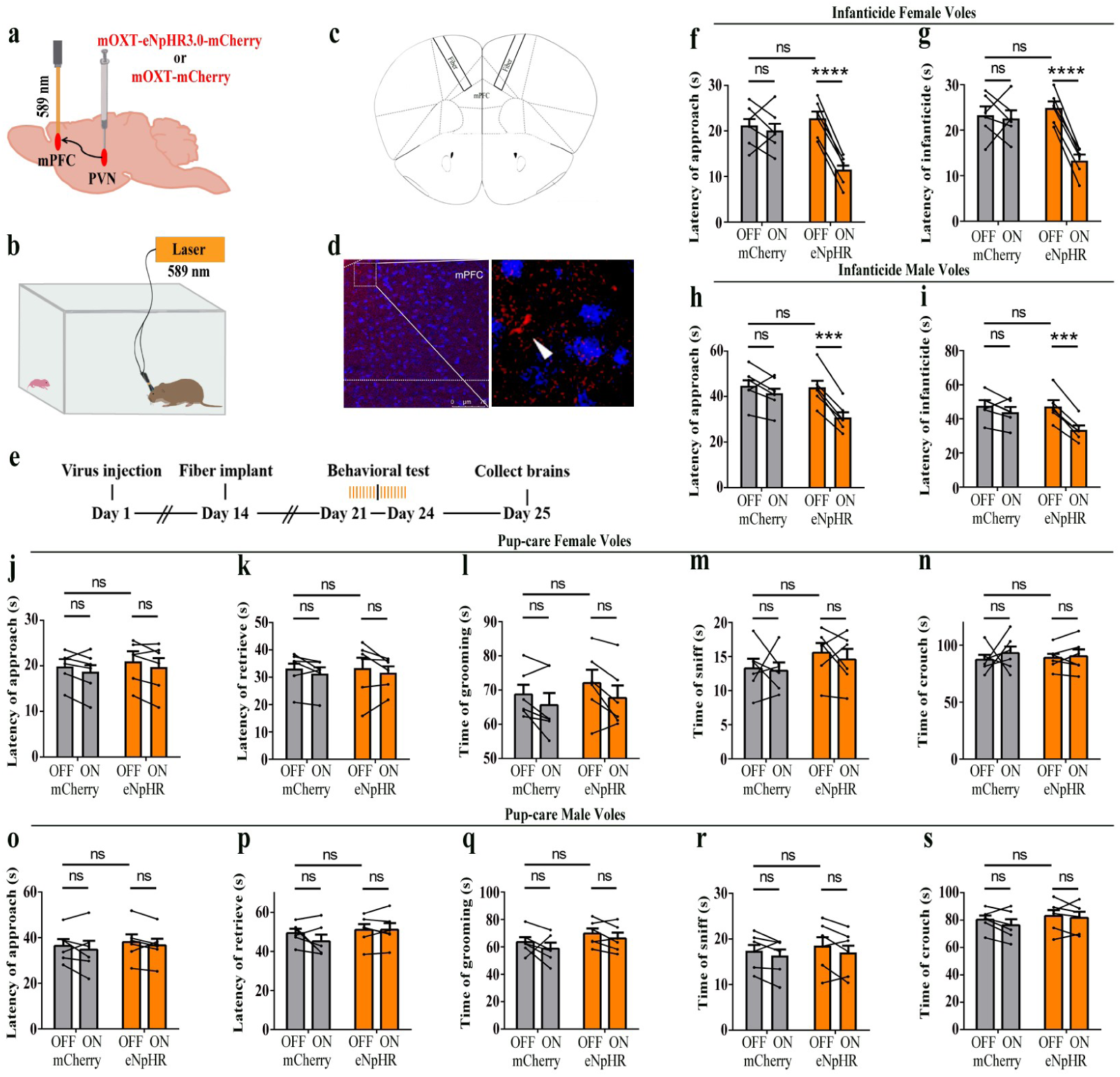
Optogenetic inhibition of the PVN OT neuron projection fibers promoted the onset of infanticide. a. Illustration of virus injection and optical fiber implantation. b. Schematic of the behavior test. c. Diagram of optical fiber implantation in the target brain region. d. Representative histological pictures of the fiber location and projection fibers. Blue, DAPI. Objective: 20x. Scale bars, 75 μm. e. Time line of the experiment. f-i, Changes in approach (f, h) and infanticide (g, i) latency of female and male infanticide voles in eNpHR (n = 6) and control mCherry groups (n = 6) before and after light deivery. \*\*\**P* < 0.001 vs. eNpHR OFF. \*\*\*\**P* < 0.0001 vs. eNpHR OFF. Two-way rmANOVA (factors: treatment × stimulus). j-n, Approach latency (j), retrieval latency (k), grooming time (l), sniffing time (m) and crouching time (n) in control mCherry (n = 6) and eNpHR (n = 6) group of pup-care female voles. o-s, Approach latency (o), retrieval latency (p), grooming time (q), sniffing time (r) and crouching time (s) in control mCherry (n = 6) and eNpHR group (n = 6) of pup-care male voles. Error bars indicate SEM. Figure 7-source data 1. Statistical results of the latency to approach and attack pups in infanticide voles, and the latency to approach and retrieve pups, duration of grooming, sniffing, and crouching in pup-care voles.

### Intraperitoneal injection of OT

For pre-clinic purpose, we tested the effect of peripheral administration of OT on pup-directed behavior (Fig. 8a). Delivery of OT promoted approach (t (4) = 3.737, *P* < 0.05, d = 1.335) and retrieval (t (4) = 4.190, *P* < 0.05, d = 2.04) in pup-care male voles (Fig. 8k, m), while it had no significant effect on the pup-directed behaviors in females (Fig. 8f-j). For infanticide voles, there was not significant prolongation of approach latency (Fig. 8d), but a significant extension of infanticide latency in males after delivery of OT (t (6) = -2.988, *P* < 0.05, d = 1.345, Fig. 8e). In infanticide female voles, both the latency to approach (Z = -2.380, *P* < 0.05, d = 5.891, Fig. 8b) and attack pups (t (7) = -3.626, *P* < 0.01, d = 1.063, Fig. 8c) were significantly prolonged by the delivery of OT. In addition, we integrated data from our pre-test of effects of OT on numbers of infanticide voles and we found that intraperitoneal injection of OT significantly reduced the number of infanticide female (χ^2^ = 4.740, *P* < 0.05, odds ratio: OR = 0.303, Fig. 8p) and male voles (χ^2^ = 3.039, *P* = 0.081, OR = 0.292, Fig. 8q). These results indicate that peripheral delivery of OT can promote the onset of pup care behavior in pup-care male voles and can significantly suppress the infanticide in both sexes. This provides a basis for the application of OT in clinic and wild life management.

**Fig. 8.**
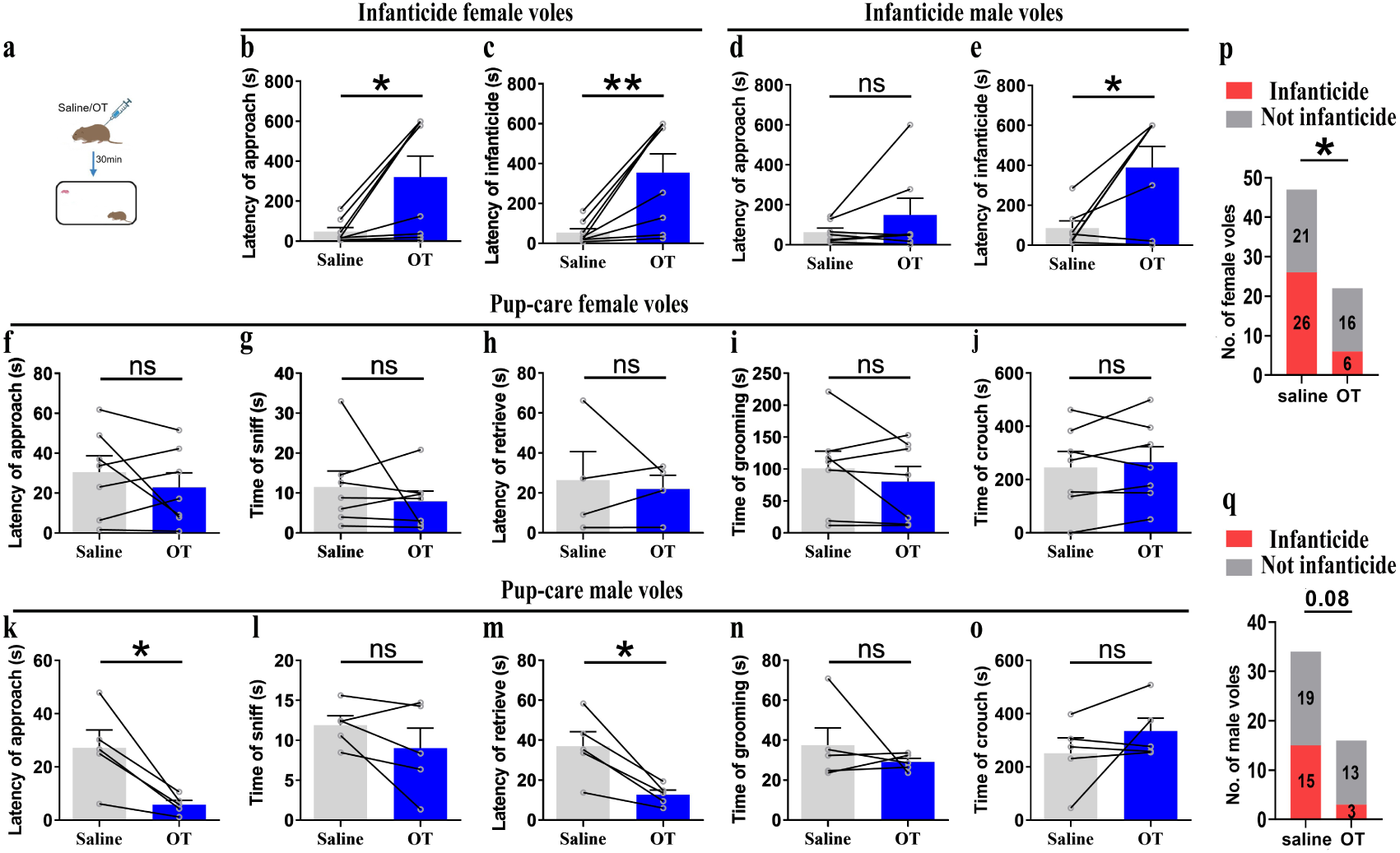
Pup-directed behaviors before and after intraperitoneal delivery of OT. a. Diagram of intraperitoneal OT delivery. b,c, Approach (b) and infanticide latency (c) of infanticide female voles (n = 8). \**P* < 0.05. Paired-samples t-test. \*\**P* < 0.01. Wilcoxon signed ranks test. d,e, Approach (d) and infanticide latency (e) of infanticide male voles (n = 7). \**P* < 0.05. Paired-samples t-test. f-j, Approach latency (f, n = 7), sniffing time (g, n = 7), latency to retrieve (h, n = 4), grooming time (i, n = 7) and crouching time (j, n = 7) before and after delivery of OT in pup-care female voles. k-o, Approach latency (k), sniffing time (l), latency to retrieve (m), grooming time (n) and crouching time (o) before and after delivery of OT in pup-care male voles (n = 5). \**P* < 0.05. Paired-samples t-test. p,q Changes in infanticide rates in female (p) and male (q) voles after administration of saline and OT. \**P* < 0.05. Pearson chi-square test. Error bars indicate SEM. Figure 8-source data 1. Statistical results of the latency to approach and attack pups in infanticide voles, and the latency to approach and retrieve pups, duration of grooming, sniffing, and crouching in pup-care voles. Statistical results on the number of infanticide or non infanticide voles induced by injection of saline and OT.

## Discussion

In this study, we used monogamous, highly social mandarin voles to explore the role of PVN OT neurons and PVN^OT^-mPFC projections in the modulation of pup care and infanticide behaviors. More OT neurons in the PVN were activated during pup care than infanticide behaviors. Optogenetic activation of the OT neurons in the PVN or OT neuron fibers in the mPFC promoted pup care and inhibited infanticide behavior, whereas inhibition of these neurons and their fibers in the mPFC promoted infanticide. In addition, intraperitoneal administration of OT promoted approach and retrieval of pups in pup care male voles, and inhibited infanticide in both male and female voles. The present study revealed that the PVN to mPFC OT neural projections regulate pup care and infanticide behavior in virgin mandarin voles.

Firstly, we found that more OT neurons in the PVN were activated during pup care than infanticide behaviors, which is consistent with the well-established prosocial role of OT and its ability to promote pup care behavior (Bosch & Young, 2018). In a previous study on virgin male prairie voles, OT and Fos colabeled neurons in PVN increased after exposure to conspecific pups and experiencing paternal care (Kenkel et al., 2012). In another study of prairie voles, OT and c-Fos colabeled neurons in the PVN significantly increased after becoming parents, which may be due to a shift from virgin to parents (Kelly, Hiura, Saunders, & Ophir, 2017). Meanwhile, we found that activation of OT neurons in the PVN facilitated pup care behaviors such as approach, retrieval and crouching in pup-care male voles; whereas inhibition of OT had no effect on paternal behavior; activation and inhibition of OT neurons in the PVN had no significant effect on pup care behaviors in pup-care females; and activation of OT neurons in the PVN inhibited pup killing in infanticide voles, whereas the corresponding inhibition of OT neurons in the PVN facilitated infanticide. This finding is consistent with previous report that silencing OT neurons delayed the retrieval behavior in virgin mice (Carcea et al., 2021). Further study found that simply observing dams retrieve pups through a transparent barrier could increase retrieval behavior and PVN OT neuron activity of virgin females (Carcea et al., 2021). If OTR knockout mice are used, no pup retrieval occurs after observation, and these results suggest that activation of PVN OT neurons in virgin mice induced by visual signals promoted pup care behaviors (Carcea et al., 2021). In addition, this study further demonstrated that OT in the PVN facilitated the retrieval behavior by modulating the plasticity of the left auditory cortex and amplifying the response of mice to the pup’s call (Carcea et al., 2021). In our another study, we found that the OT neurons in the PVN projecting to the VTA as well as to the Nac brain region regulate pup-directed behaviors, which may also be accompanied by dopamine release (He et al., 2021). This studies also support finding from optogenetic activation of OT neurons in the PVN in the present study. The results of the present study are also supported by a recent study that chemogenetic activation OT neurons in the PVN increase pup care and reduce infanticide (Inada et al., 2022).

However, manipulation of OT neurons in the PVN produced no significant effect on pup caring in pup-care females. This may be due to the fact that female voles have inherently higher OT neural activity (Häussler, Jirikowski, & Caldwell, 1990), and female mice have more OT neurons and OT axon projections than males (Häussler et al., 1990), and that there are also significant differences in OTR expression between two sexes (Insel, Gelhard, & Shapiro, 1991; Uhl-Bronner, Waltisperger, Martínez-Lorenzana, Condes Lara, & Freund-Mercier, 2005) that possibly shows ceiling effects of the OT system. In the present study, we found that females have more activated OT neurons (Figure 1, d, g) and released higher levels of OT into the mPFC (Figure 4 d, e) than males. This sex difference has been reported in other study that activation of OT neurons in the PVN activated noradrenergic neurons in the locus coeruleus by co-releasing OT and glutamate, increased attention to novel objects in male rats, and that this neurotransmission was greater in males than in females (Wang, Escobar, & Mendelowitz, 2021). In a study on virgin female mice, pup exposure was found to activate oxytocin and oxytocin receptor expressing neurons (Okabe et al., 2017). Virgin female mice repeatedly exposed to pups showed shorter retrieval latencies and greater c-Fos expression in the preoptic area (POA), concentrations of OT in the POA were also significantly increased, and the facilitation of alloparental behavior by repeated exposure to pups occurred through the organization of the OT system (Okabe et al., 2017). In the present study, we also observed that optogenetic activation of OT neurons in PVN increased crouching behavior in the pup-care male voles, but did not affect grooming time possibly via increase of OT release. This result is in line with a previous study that injection of an OTR antagonist into the MPOA of male voles significantly reduced the total duration of pup care behavior and increased the latency to approach pups and initiate paternal behavior in male voles (Yuan et al., 2019). This finding suggests that the results of peripherally administered OT and optogenetically activated PVN OT neurons in the present study may also have the involvement of OT-OTR interactions in MPOA. Parturition in experimental animals is accompanied by a decrease in infanticide and the emergence of pup-care behavior, and along with this process OTR expression increased not only on mammary contractile cells, but also in various regions of the brain, such as MPOA, VTA, and OB, which are all considered to be important brain regions related to the onset and maintenance of pup care behaviors (Bosch & Neumann, 2012; Shahrokh, Zhang, Diorio, Gratton, & Meaney, 2010; Yu, Kaba, Okutani, Takahashi, & Higuchi, 1996). For example, in a OB study, intra-OB injection of OT antagonist significantly delayed the onset of pup care behaviors such as retrieving pups, crouching, and nesting in female rats, whereas intra-OB injection of OT in virgin females induced 50% of females to show intact pup care behaviors (Yu et al., 1996). Therefore, the effects of activation of PVN OT neurons may be a result of actions on multiple brain regions involved in the expression of pup care behavior. Which brain regions that PVN OT neurons project to are involved in pups caring or infanticide needs further studies. Although we used a virus strategy to specifically activate or inhibit PVN OT neurons, other neurochemical may also be released during optogenetic manipulations because OT neurons may also release other neurochemicals. In one of our previous studies, activation of the OT neuron projections from the PVN the VTA as well as to the Nac brain also altered pup-directed behaviors, which may also be accompanied by dopamine release (He et al., 2021). In addition, back-propagation of action potentials during optogenetic manipulations may also causes the same behavioral effect as direct stimulation of PVN OT cells. These indirect effects on pup-directed behaviors should also be investigated further in the future study.

Optogenetic activation of the OT neural projection fibers from the PVN to the mPFC facilitated the onset of pup care behaviors, such as approach and retrieval, in pup-care male voles, whereas inhibition of this circuit had no effect; neither activation nor inhibition had a significant effect in pup-care females; and activation of this neural circuit inhibited infanticide in infanticide voles, whereas the corresponding inhibition of this circuit facilitated infanticide. In addition, we have demonstrated an increase in OT release in pup-care voles upon approaching and retrieving pups, and a decrease in OT release in infanticide voles when infanticide occurs, as recorded by OT sensors in the mPFC. There is some evidence indicating that the mPFC may be an important brain region for OT to exert its effects. In addition to expressing OTRs in the mPFC (W. Liu, Pappas, & Carter, 2005; Smeltzer et al., 2006), the mPFC contains OT-sensitive neurons (Ninan, 2011) and receives projections of OT neurons from the hypothalamus (Knobloch et al., 2012; Sofroniew, 1983). It has been further shown that blocking OTR in the mPFC of postnatal rats by an OTR antagonist delayed the retrieval of pups and reduced the number of pups retrieved by rats, impaired the care of pups, and decreased the latency to attack intruders, increased the number of attacks, and increased anxiety in postnatal rats but had no effect on the level of anxiety in virgin rats (Sabihi, Dong, Durosko, & Leuner, 2014). These results further suggest that OT in the mPFC is involved in the regulation of pup care behavior and support the results of this study in pup care voles and infanticide voles from an OTR perspective. Previous studies have shown that optical activation of the mPFC maintains aggression within an appropriate range, with activation of this brain region suppressing aggression between male mice and inhibition of the mPFC resulted in quantitative and qualitative escalation of aggression (Takahashi, Nagayasu, Nishitani, Kaneko, & Koide, 2014), which is similar to our findings with infanticide voles. For pup care behavior it has been reported that the mPFC brain region may be involved in the rapid initiation of pup care behavior in mice without pairing experience (Alsina-Llanes & Olazábal, 2020), which supported the experimental results of pup-care male voles in the present study, but the present study further suggested that OT neurons projecting to mPFC regulated pup-caring and infanticide behavior possibly via increase of OT release in the mPFC. We must also mention the function of the mPFC subregion. In the present study, virus were injected into the PrL. The PrL and IL regions of the mPFC play different roles in different social interaction contexts (Bravo-Rivera, Roman-Ortiz, Brignoni-Perez, Sotres-Bayon, & Quirk, 2014; Moscarello & LeDoux, 2013). A study has shown that the PrL region of the mPFC contributes to active avoidance in situations where conflict needs to be mitigated, but also contributes to the retention of conflict responses for reward (Capuzzo & Floresco, 2020). This may reveal that the suppression of infanticide by PVN to mPFC OT projections is a behavioral consequence of active conflict avoidance. In a study of pain in rats, OT projections from the PVN to the PrL were found to increase the responsiveness of cell populations in the PrL, suggesting that OT may act by altering the local excitation-inhibition (E/I) balance in the PrL (Y. Liu et al., 2023). A study of anxiety-related behaviors in male rats suggests that the anxiolytic effects of OT in the mPFC are PL-specific and that this is achieved primarily through the engagement of GABAergic neurons, which ultimately modulate downstream anxiety-related brain regions, including the amygdala (Sabihi, Dong, Maurer, Post, & Leuner, 2017). This may provide possible downstream pathways for further research.

Another interesting finding is that peripheral OT administration promoted pup care behavior in male voles, such as approaching and retrieving pups, but had no significant effect on pup care behaviors in female voles. This result was supported by previous study that OT administered peripherally inhibited infanticide in pregnant and reproductively inexperienced females and promoted pup caring in pregnant females (McCarthy, Bare, & vom Saal, 1986). Further research has demonstrated that OT in the central nervous system also inhibited infanticide in female house mice (McCarthy, 1990). In addition, peripheral administration of OT inhibits infanticide in male mice without pairing experience (Nakahara et al., 2020). Moreover, the experience of pairing facilitated the retrieval of pups by increasing peripheral OT levels in both male and female mice (Lopatina et al., 2011). This is similar to the findings of pup-care males in the present study, whereas the absence of similar results in females may be due to differences in OT levels between the sexes (Tamborski, Mintz, & Caldwell, 2016). Pup-care female voles in this study already showed short latency to approach and retrieve pups before peripheral administration of OT. We also found that peripheral OT administration suppressed infanticide behavior in both male and female voles. It is consistent with previous study that infanticide in female house mice with no pairing experience and pregnant females with pre-existing infanticide was effectively suppressed by subcutaneous injection of OT, while injection of OT also helped pregnant females to show care for strange pups (McCarthy et al., 1986). Unpaired male mice’s aggression towards pups was accompanied by changes in the activity of the vomeronasal neurons, and as males were paired with females and lived together, the activity of these neurons decreased, accompanied by a shift from infanticide to pup care behavior (Nakahara et al., 2020). The OTR expresses throughout the vomeronasal epithelium that allows OT to potentially inhibit infanticide by reducing vomeronasal activity. The previous study found that intraperitoneal injection of OT reduced the activity of the vomeronasal, and further validated that OT modulates the activity of vomeronasal neurons by acting on the sensory epithelium to produce behavioral changes by intraperitoneal injection of an OTR antagonist that cannot cross the blood-brain barrier (Nakahara et al., 2020). It has recently been shown that peripheral OT was able to cross the blood-brain barrier into the central nervous system to act (Yamamoto et al., 2019), meaning that delivery of OT via the periphery may have increased central OT levels and thus exerted an effect. A recent study demonstrated that the secretion of OT in the brain of male mice with no pairing experience facilitated the performance of pup care behaviors and inhibited infanticide (Inada et al., 2022), which also supported the results of the present study. Similar to rodents and non-human primates, there is evidence to suggest that OT contributes to paternal care (Feldman & Bakermans-Kranenburg, 2017). Fathers with partners have higher plasma OT levels than non-fathers without partners (Mascaro, Hackett, & Rilling, 2014). Intranasal OT treatment increases fathers’ play, touch, and social interaction with their children (Weisman, Zagoory-Sharon, & Feldman, 2012). Our result provide possible application of OT in reduction of abnormality in pup-direct behavior associated with psychological diseases in human such as depression and psychosis (Milia & Noonan, 2022; Naviaux, Janne, & Gourdin, 2020) and increases of well being of wild life.

In summary, these results indicate that the PVN to mPFC OT neural projection is involved in the regulation of pup care and infanticide behavior in virgin mandarin voles. These data provide new insights into the neural circuits underlying OT-mediated pup-directed behaviors.

## Supporting information

revised source data

---

**Supplementary Data Fig. 1.**
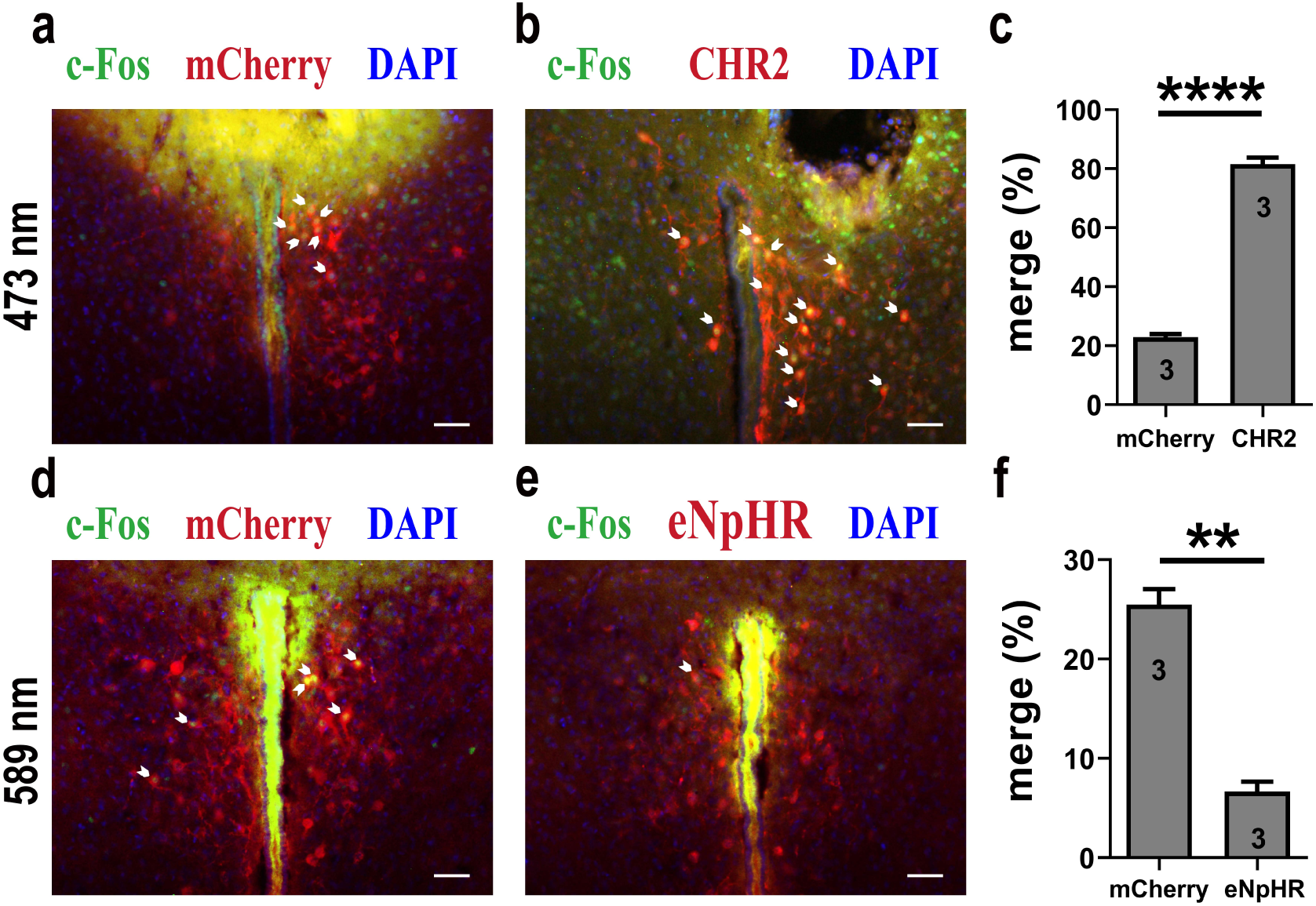
Light-induced c-Fos expression overlapping with mCherry, CHR2 or eNpHR. a, 473 nm light-induced c-Fos (green) expression overlapping with mCherry (red). Blue, DAPI. Objective: 20x. Scale bars, 50 μm. b. 473 nm light-induced c-Fos (green) expression overlapping with CHR2 (red). Blue, DAPI. Objective: 20x. Scale bars, 50 μm. c, 473 nm light induced more c-fos expression in CHR2, which indicated the effectiveness of CHR2 (n = 3). \*\*\*\**P* < 0.0001. Independent-samples t-test. d, 589 nm light-induced c-Fos (green) expression overlapping with mCherry (red). Blue, DAPI. Objective: 20x. Scale bars, 50 μm. e, 589 nm light-induced c-Fos (green) expression overlapping with eNpHR (red). Blue, DAPI. Objective: 20x. Scale bars, 50 μm. f, 589 nm light induced less c-fos expression in eNpHR, which indicated the effectiveness of eNpHR (n = 3). \*\**P* < 0.01. Independent-samples t-test. Error bars indicate SEM. For c,f, n = 3 per group, data are mean ± s.e.m. Statistical analysis was performed using independent-samples T test (c,f). \*\**P* < 0.01, \*\*\*\**P* < 0.0001. Details of the statistical analyses are as follows: c, t (4) = -23.139, *P* = 0.000021, f, t (4) = 10.136, *P* = 0.001. Supplementary Data Fig. 1-source data 1 Statistical results of light-induced c-Fos expression overlapping with mCherry, CHR2 or eNpHR.

**Supplementary Data Fig. 2.**
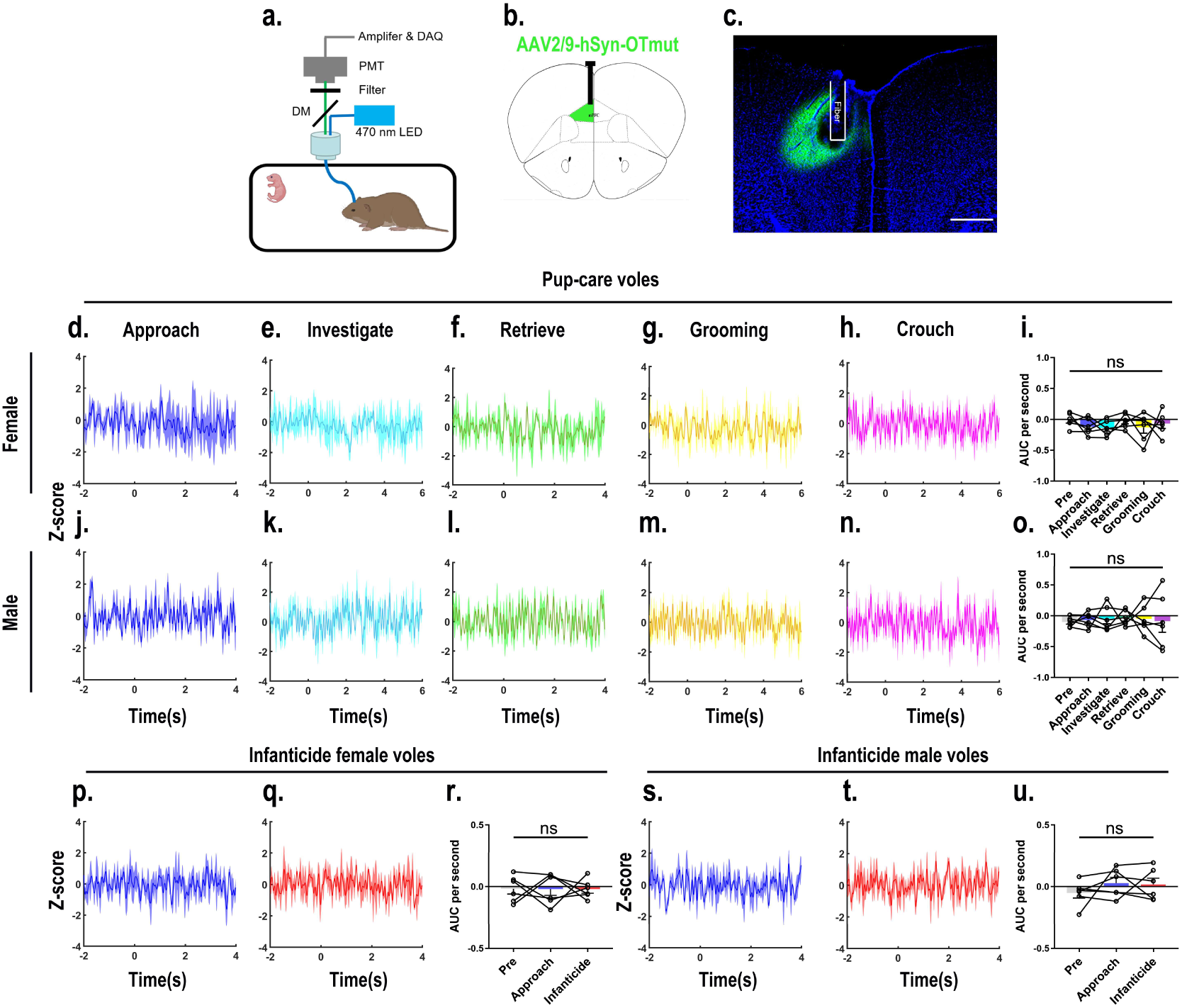
Recordings of OTmut sensor signals in the mPFC on OT release. a. Recording instrument settings. b. Illustrations of viral expression and optical fibre location. c. Representative histological image of OTmut sensor (green) and optical fibre locations. Blue, DAPI. Objective: 4x. Scale bars, 500 μm. d-h,j-n, PETHs of z-score of OTmut sensor signals for the following pup-directed behaviors: approach (d,j), investigate (e,k), retrieve (f,l), grooming (g,m) and crouch (h,n). i,o, AUC per second of z-scores for pup-care female (n = 6, i) and male (n = 6, o) voles across various pup-directed behaviors. p,q,s,t, PETHs of z-score of OTmut sensor signals for approach and infanticide in infanticide voles. r,u, AUC per second of z-score of OTmut sensor signals for pre-pup exposure, approach and infanticide in infanticide female (n = 6, r) and male (n = 6, u) voles. Error bars indicate SEM. For i,o,r,u, n = 6 per group, data are mean ± s.e.m. Statistical analysis was performed using one-way rmANOVA (i,o,r and u). Details of the statistical analyses are as follows: i, F (5, 25) = 0.680, *P* = 0.643; o, F (2, 25) = 0.074, *P* = 0.996; r, F (2, 10) = 0.002, *P* = 0.998; u, F (2, 10) = 0.665, *P* = 0.536. Supplementary Data Fig. 2-source data 1 Area under the curve per second for pre-pup exposure, approach, and infanticide behaviors in infanticide voles and area under the curve per second for pre-pup exposure, approach, investigation, retrieval, grooming, and crouching behaviors in pup-care voles.

**Supplementary Data Fig. 3.**
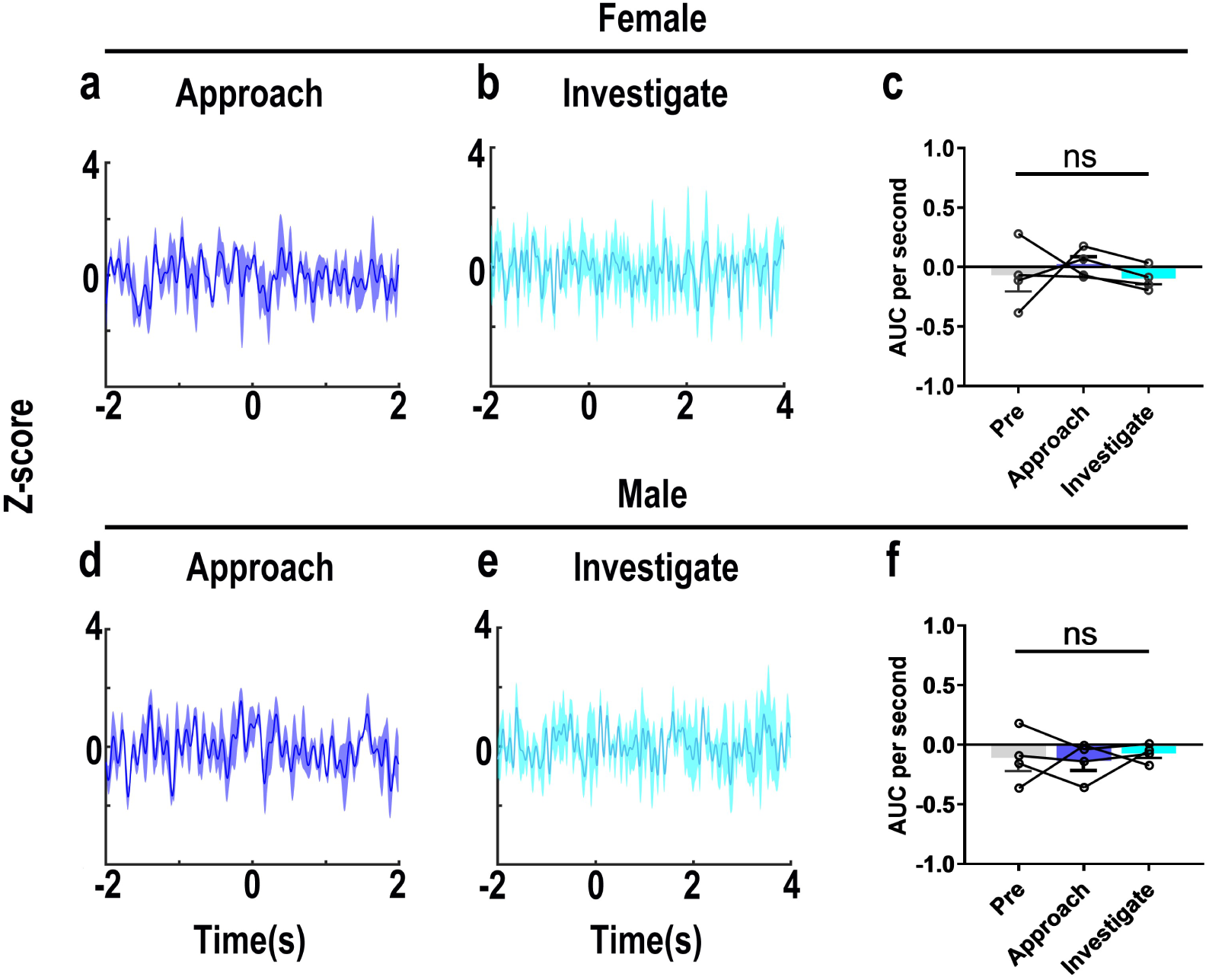
Recordings of OT1.0 sensor signals for investigating object. a,b,d,e, Post-event histograms (PETHs) of z-score of OT1.0 sensor for approaching and investigating object in female (a,b) and male (d,e) voles. c,f, AUC per second of z-scores for pre object, approaching and investigating behaviors in female (n = 4, c) and male (n = 4, f) voles. Error bars indicate SEM. For c,f, n = 4 per group, data are mean ± s.e.m. Statistical analysis was performed using one-way rmANOVA. Details of the statistical analyses are as follows: c, F (1.016, 3.047) = 0.346, *P* = 0.601; f, F (2,6) = 0.183, *P* = 0.838. Supplementary Data Fig. 3-source data 1 Area under the curve per second for pre-object exposure, approach, and investigating in female and male voles.

**Supplementary Data Fig. 4.**
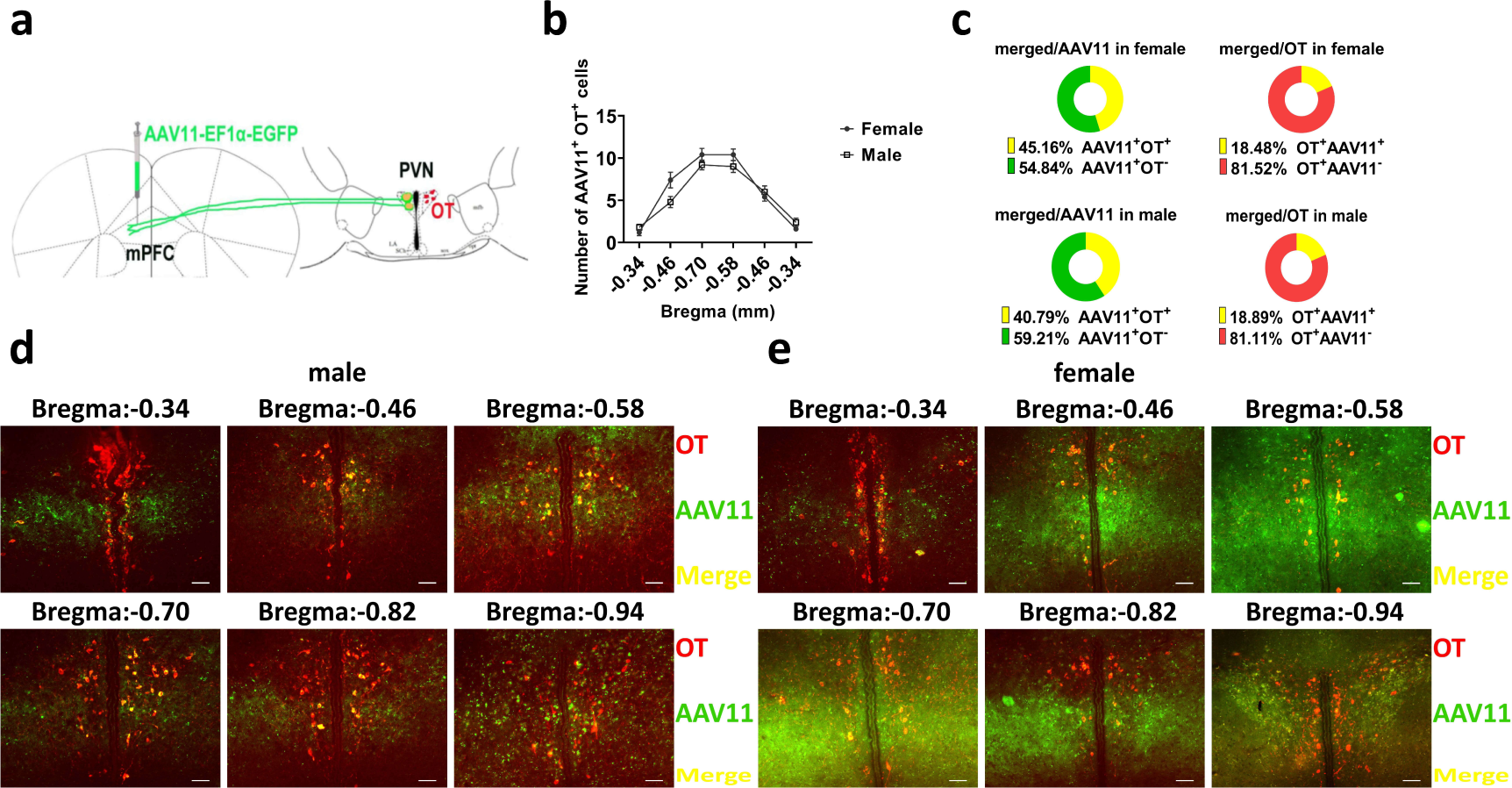
Number of PVN OT neurons projecting to mPFC. a. Schematic diagram of virus injection and immunohistological staining. b. Statistics of PVN to mPFC OT projection extent. c. Percentage of OT positive cells in PVN projecting to mPFC and percentage of OT cells in PVN that project to mPFC. d. e. Representative histologic images of male and female. Red, OT. Green, AAV11. Yellow, merged. Objective: 20x. Scale bars, 50 μm. Supplementary Data Fig. 4-source data 1 Counts of AAV11, OT and co-expressed cells.

**Supplementary Data Fig. 5.**
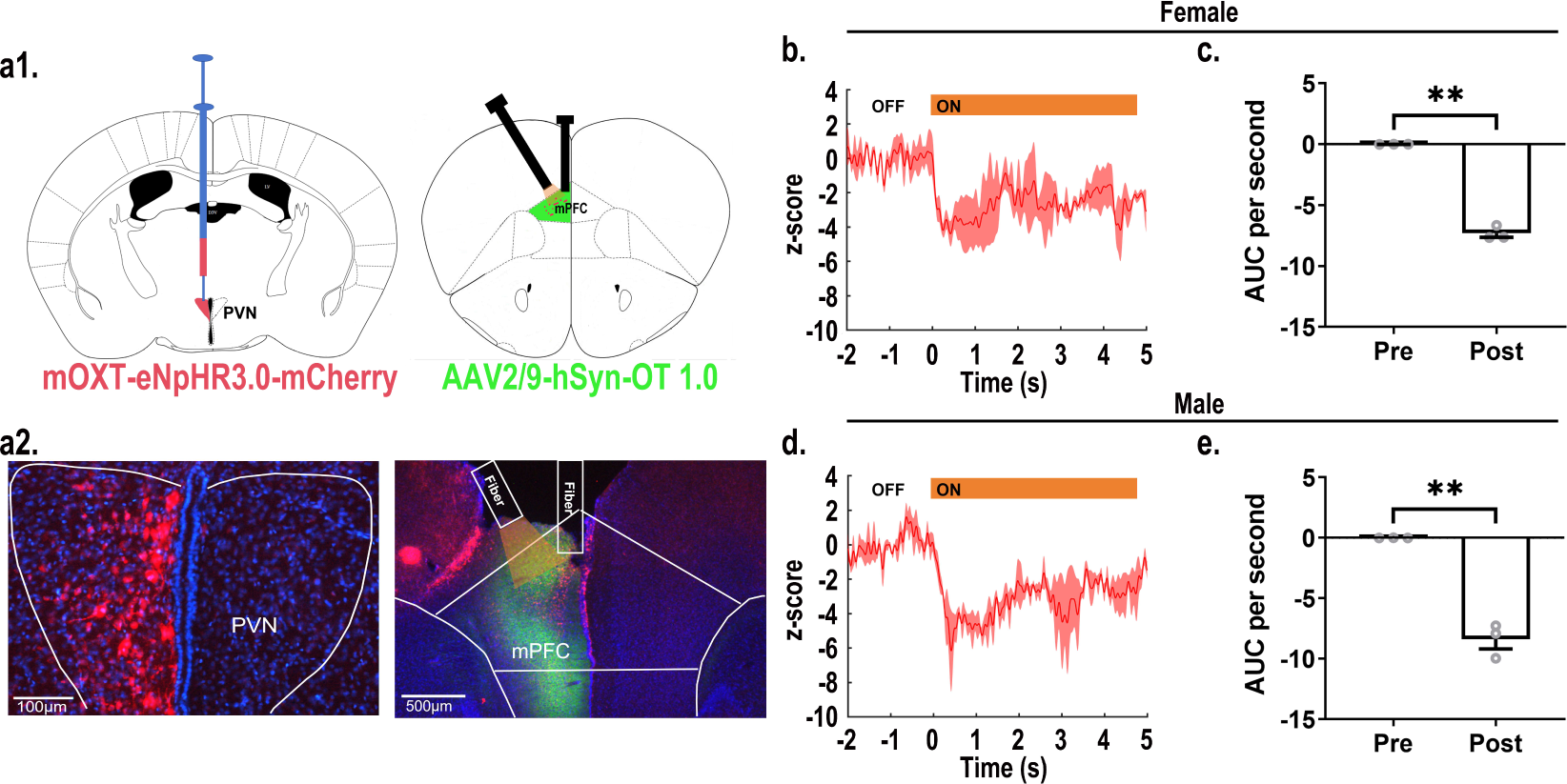
Recordings of OT release during optogenetic inhibition of OT fibers in the mPFC. a1. Schematic diagram of virus injection and optical fiber implanting. a2. Representative histological images of viruses infection and fibers placement. Left: Red: eNpHR3.0, Blue: DAPI, objective: 20x, scale bars: 100 μm; Right: Red: eNpHR3.0, Green: OT1.0 sensors, Blue: DAPI, objective: 4x, scale bars: 500 μm. b. Representative histogram of OT release in female at 589 nm light. c. AUC per second of OT1.0 sensors pre and post inhibition of optogenetics in females (n = 3). \*\**P* < 0.01, Paired-samples t-test. d. Representative histogram of OT release in male at 589 nm light. e. AUC per second of OT release pre and post optogenetic inhibition in males (n = 3). \*\**P* < 0.01, Paired-samples t-test. Supplementary Data Fig. 5-source data 1 Statistics of AUC before and after optogenetic inhibition.

**Supplementary Data Fig. 6.**
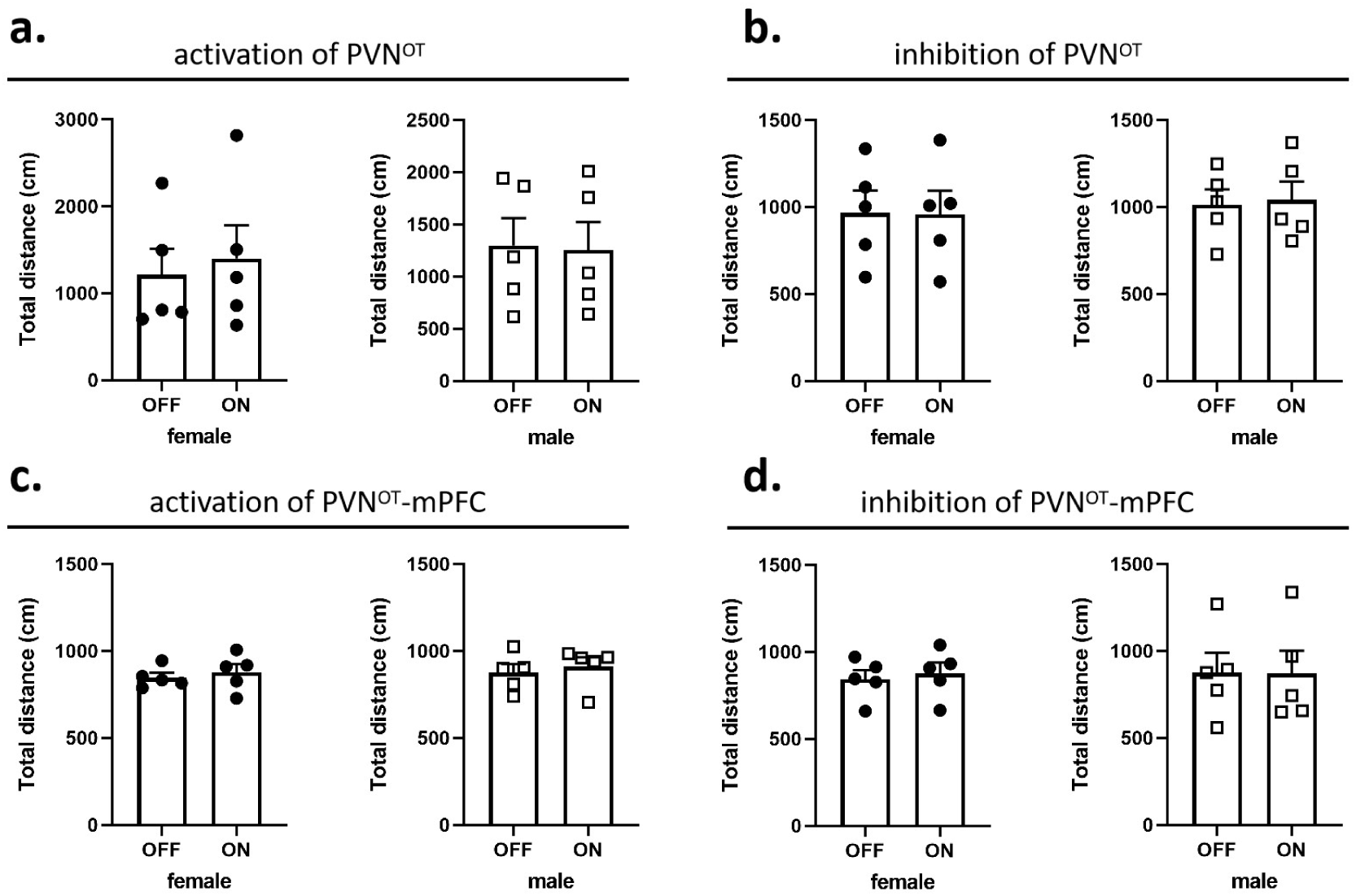
Effects of optogenetic manipulations on locomotion of the subjects. a. b. Total distance traveled by the subject before and after activation or inhibition of the PVN OT neurons. c. d. Total distance traveled by the subject before and after activation or inhibition of the PVN OT neurons projections to the mPFC. Supplementary Data Fig. 6-source data 1 Statistics of distance traveled before and after optogenetic manipulation.

**Supplementary Data Fig. 7.**
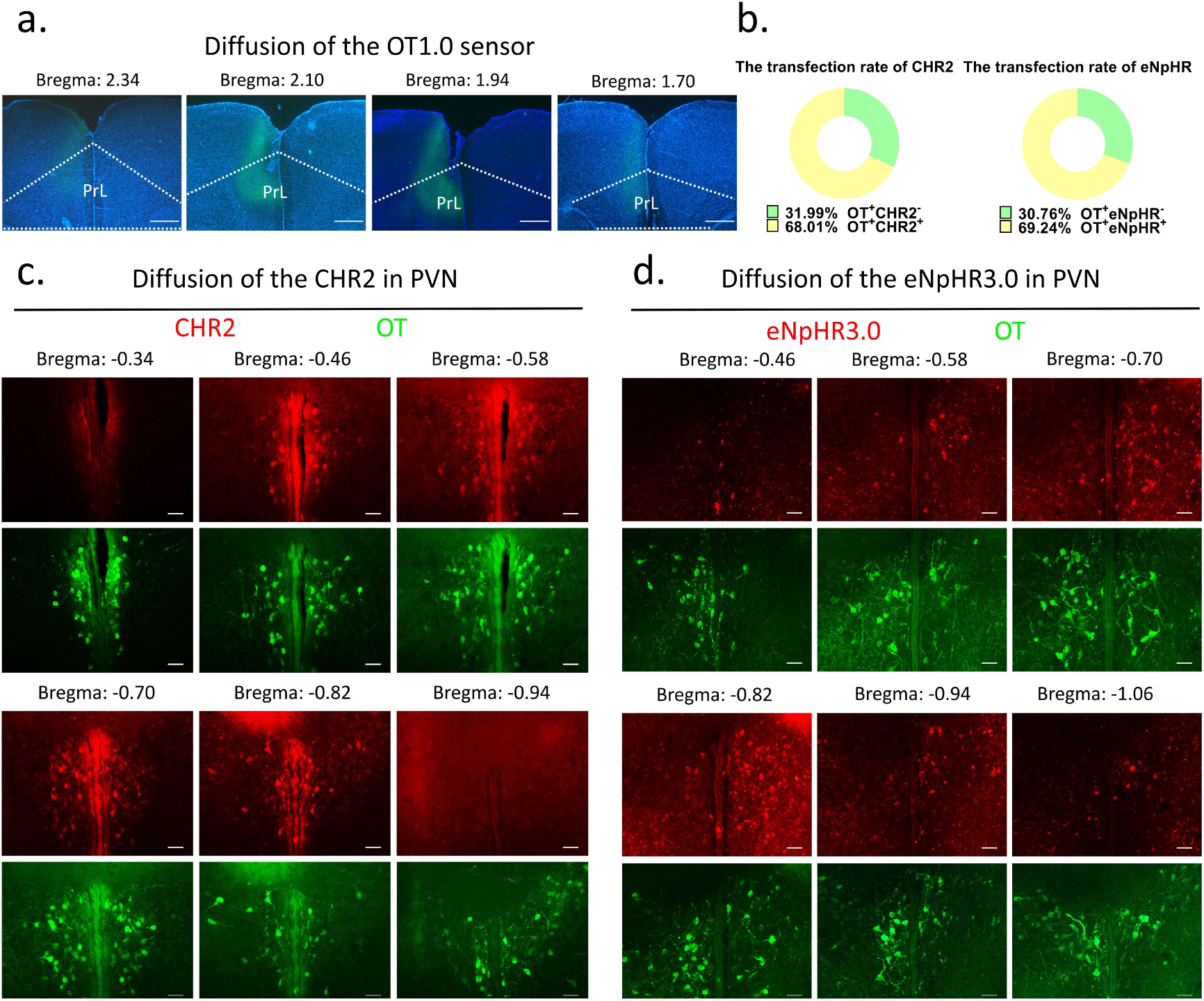
Diffusion of centrally injected agents. a. Diffusion of OT1.0 sensor in PrL. Green: OT1.0 sensors, Blue: DAPI, Objective: 4x. Scale bars: 500 μm. b. The transfection rate of chr2 and eNpHR3.0 in OT cells. c. Diffusion of CHR2 in PVN. Green: OT, Red: CHR2, Objective: 20x. Scale bars: 50 μm. d. Diffusion of eNpHR3.0 in PVN. Green: OT, Red: eNpHR3.0, Objective: 20x. Scale bars: 50 μm. Supplementary Data Fig. 7-source data 1 Counting of CHR2 and eNpHR3.0 virus-transfected OT cells.

## Methods

### Animals

Mandarin voles were captured from the wild in Henan, China. The experimental protocols were approved by the Animal Care and Use Committee of Shaanxi Normal University, and was conducted in accordance with the ethical principles of animal use and care of China. The virgin Mandarin voles (*Microtus mandarinus*) used in this study were F2 generations that were bred in Animal Center of Shaanxi Normal University and were kept at 24℃ under a 12h light-dark cycle (lights on at 8 a.m.) with food and water provided ad libitum. Before the experiments, we expose the animals to pups, and subjects may exhibit pup care, infanticide, or neglect; we group subjects according to their behavioral responses to pups, and individuals who neglect pups are excluded. The stereotactic surgery was performed at the age of 8 weeks of voles. After surgery, voles were housed with their cage mates. Behavioral tests were carried out 3weeks after surgery for animal recovery and the viral infection, and 1-5 days old pups were from other breeders. In case the pups were attacked, we removed them immediately to avoid unnecessary injuries, and injured pups were euthanized.

### Viruses

AAV2/9-mOXT: Promoter-hCHR2(H134R)–mCherry-ER2-WPRE-pA (8.41×10^12^ μg/ml) and AAV2/9-mOXT: Promoter-mCherry-pA (1.21×10^13^ μg/ml) were purchased from Shanghai Taitool Bioscience LTD. AAV2/9-mOXT-eNpHR3.0-mCherry-WPRE-hGH-pA (2.27×10^12^ μg/ml) were purchased from BrainVTA (Wuhan, China) LTD. AAV2/9-hSyn-OT 1.0 (2.06×10^12^ μg/ml), AAV2/9-hSyn-OTmut (2.10×10^12^ μg/ml) and AAV11-EF1α-EGFP (5.00×10^12^ μg/ml) were purchased from Brain Case Biotechnology LTD. The details about the construction of CHR2 and mCherry viruses used in optogenetic manipulation can refer to a previous study in which they constructed an rAAV-expressing Venus from a 2.6 kb region upstream of OT exon 1, which is conserved in mammalian species (Knobloch et al., 2012). The details about construction of the eNpHR 3.0 virus can refer to one study in which expression of the vector is driven by the mouse OXT promoter, a 1kb promoter upstream of exon 1 of the OXT gene, which has been shown to induce cell type-specific expression in OXT cells (Peñagarikano et al., 2015). Details about the construction of OT 1.0 sensor can be referred to the research of Professor Li’s group (Qian et al., 2023). All viruses were dispensed and stored at -80℃.

### Immunohistochemistry

After behavioral tests, serial brain sections were harvested for histological analysis. Anesthetized voles were perfused with 40ml of PBS and 20ml of 4% paraformaldehyde. After perfusion, brains were excised and post-fixed by immersion in 4% paraformaldehyde overnight at 4℃. Brains were dehydrated in 20% and then 30% sucrose for 24h respectively before they were embedded in OCT and cryosectioned into 40 μm slices. Brain slices were rinsed with PBS (10 min) and PBST (PBS and 0.1% Triton X-100, 20 min), blocked in ready-to-use goat serum (Boster, AR0009) for 30 minutes at room temperature (RT), and then incubated overnight at 4℃ with primary antibody in PBST. The primary antibodies used were: mouse anti-OT (1:7000, Millipore, MAB5296), rabbit anti-c-Fos (1:1000, Abcam, ab190289). Following PBST washing (3×5 min), sections were incubated with secondary antibodies in PBST for 2h (RT) and stained with DAPI, and then washed once more with PBS (3×5 min). The secondary antibodies used were: goat anti-mouse Alexa Fluor 488 (1:200, Jackson ImmunoResearch, 115-545-062), goat anti-rabbit Alexa Fluor 488 (1:200, Jackson ImmunoResearch, 111-545-003), goat anti-rabbit TRITC(1:200, Jackson ImmunoResearch, 111-025-003) and ready-to-use DAPI staining solution (Boster, AR1177). Finally, the brain slices were sealed with an anti-fluorescent attenuation sealer.

Images were captured with a fluorescent microscope (Nikon) to confirm the viral expression, placements of optic fiber and viruses, and also the number of c-Fos, OT and virus positive cells. To analyze the activity of OT neurons (co-expression of c-Fos and OT) among different behaviors and the specificity of viral expression (co-expression of viruses and OT) in the PVN brain region, Brain slices of 40 µm were collected consecutively on 4 slides, each slide had 6 brain slices spaced 160 µm apart from each other, and counting was performed on one of the slides. Positive cells in the PVN were manually counted based on the Allen Mouse Brain Atlas and our previous studies.

### Stereotaxic Surgery

For optogenetic manipulation experiments, CHR2, eNpHR and mCherry-expressing control virus were stereoaxically injected into the PVN (AP: -0.4 mm, ML: 0.2 mm, DV: 5.3mm) bilaterally through a Hamilton needle using nanoinjector (Reward Life Technology, KDS LEGATO 130) at 100 nl/min. For optogenetic manipulation, optic fibers (2.5 mm O.D., Reward Life Technology, China) with an appropriate fiber length (PVN: 6 mm, mPFC: 3 mm) were implanted ∼100 μm above the PVN and mPFC (AP: 2.2 mm ML: 0.9, DV: 1.89 with a 20° angle lateral to middle) and secured with dental cement (Changshu ShangChi Dental Materials Co., Ltd.,202005). For fiber optometry experiment, optic fibers were inserted ∼100 μm above the mPFC (AP: 2.2 mm ML: 0.3, DV: 1.8) after injecting the OT1.0 sensor viruses. The stereotaxic coordinates were determined from three-dimensional brain atlas (Chan, Kovacevíc, Ho, Henkelman, & Henderson, 2007) and the adjusted coordinates in our lab. The stereotaxic coordinates were determined by the Allen Mouse Brain Atlas and laboratory-corrected data used for voles. Individuals with appropriate viral expression and optical fiber embedding location were included in the statistical analysis, otherwise excluded. The diffusion of central optogenetic viruses and OT1.0 sensors are shown in the supplemental figure (Supplementary Data Fig. 7).

### Optogenetic

Test animals were injected with 300 nL AAV2/9-mOXT: Promoter-hCHR2(H134R)–mCherry-ER2-WPRE-pA or rAAV-mOXT-eNpHR3.0-mCherry-WPRE-hGH-pA bilaterally into the PVN at 100 nl/min. Control animals were injected with AAV2/9-mOXT: Promoter-mCherry-pA in the same condition. Two weeks later, an optic fiber was implanted ∼100 μm above the PVN and mPFC bilaterally and was secured using dental cement. After surgery, animals were housed with their cage mates as before. Before the behavioral test, each implanted fiber was connected to a light laser (NewdoonInc., Aurora 300, 473 nm for activation, 589 nm for inhibition) with a 400 μm patch cord and then we introduced the test animal into center of test arena. After the test animal acclimated to the arena for at least 20 min and settled down, we placed a pup in the farthest corner away from the test animal and immediately began recording and applying light stimulation.

In pup-directed pup care behavioral test, light stimulation lasted for 11 min. Parameters used in optogenetic manipulation of PVN OT neurons were ∼ 3 mW, 20 Hz, 20 ms, 8 s ON and 2 s OFF and parameters used in optogenetic manipulation of PVN OT neurons projecting to mPFC were ∼ 10 mW, 20 Hz, 20 ms, 8 s ON and 2 s OFF to cover the entire interaction. In the infanticide behavioral test, the stimulation lasted until the pup was removed. Each vole was tested twice successively, more than 30 min apart, once with the stimulation OFF, once ON. The effect of optogenetic manipulation induced locomotion on behavioral responses to pups was exluded by recording the total distance traveled by voles without and with light stimulation for 5 minutes, respectively (Supplementary Data Fig. 6).

To confirm that ChR2 or eNpHR3.0 stimulation indeed induced neural activation or inhibition, we used light to stimulate the brain through an optical fiber when voles were alone in their home cage, and subsequently determined neural activation or inhibition by c-Fos staining one and a half hours after light stimulation.

### Fiber photometry

To record the fluorescence signals of the OT1.0 sensor during various pup-directed behaviors. Virgin voles were anesthetized with 1-2% isoflurane and immobilized on a stereotaxic device (RWD, China). Then, 300 nl AAV9-hSyn-OT1.0 or AAV9-hSyn-OTmut virus was injected into the left side of mPFC (AP: 2.2 mm ML: 0.3, DV: 1.8). After the injection, a 200 μm optical fiber was implanted ∼100 μm above the injection site and fixed with dental cement. After 2 weeks of recovery, the optic fiber was connected to the fiber photometry system (QAXK-FPS-LED, ThinkerTech, Nanjing, China) through a patch cable. This system can automatically exclude motion artifacts by simultaneously recording signals stimulated by a 405 nm light source. To avoid bleaching the sensor, the 470 nm laser power at the tip of the fiber was adjusted to 50 μW. Voles were placed in test cages and allowed to move freely for at least 20 minutes to acclimate to the environment. Then, a pup was placed in the cage at a distance from the testing vole. If the vole ignored the pup completely or attacked the pup, gently removed the pup and introduced another pup ∼60 seconds later to stimulate more interaction. This process was repeated 3-6 times, and then the vole exhibiting pup care behavior was allowed to freely interact with the last introduced pup until the vole crouched over the pup for more than 10 s, after which the pups were removed for about 60 s and reintroduced. This latter process was repeated 3-4 times. After the pup test, we subsequently placed an object (a vole-sized plastic toy) into the cage and recorded 4 voles that investigated the object 3-6 times.

Videos were recorded with screen-recoding software to synchronize the OT1.0 fluorescence signals and pup-directed behaviors. Fluorescence signals were recorded into MATLAB mat files and analyzed with customized MATLAB code. Data were matched to a variety of behaviors towards pups based on individual trials. The change in signal was displayed as z-scored ΔF/F, which was measured by (V_signal_ - mean (V_basal_))/std (V_basal_). The V_signa_ and V_basal_ refer to the recorded values at each time point and the recorded values during the baseline period before the stimuli. The AUC (area under the curve) was calculated based on z-scored ΔF/F matching the duration of the behavior, and the AUC per second was used to compare the different fluorescence signals of behaviors and the baseline.

For the combination of optogenetic inhibition and fiber optometry experiment, optogenetic virus and OT1.0 sensor were injected as described above, optic fibers were inserted above the mPFC (OT1.0 fibers: AP: 2.2 mm ML: 0.2, DV: 1.7; optogenetic fibers: AP: 2.2 mm ML: 2.0, DV: 2.4 with a 45° angle lateral to middle). The signals of OT1.0 sensor were recorded while neurons were optogenetically inhibited.

### Behavioral paradigm and analysis

Animal behaviors in optogenetic experiments were recorded by a camera from the side of a transparent cage. ‘Approach pup’ was defined as the testing vole faced and walked right up to pup, and the latency to approach was the period from the time the pup was placed in the cage until the vole began to approach the pup. ‘Investigate pup’ was defined as the vole’s nose came into close contact with any part of the pup’s body. ‘Attack pup’ was defined as the vole attacked or bit a pup which can be recognized by the wound, and the latency to attack was the period from the time the pup was placed in the cage until the vole launched an attack. ‘Retrieve pup’ was defined as from the time a vole picked up a pup using its jaws to the time it dropped the pup at or around the nest, and the latency of retrieval was the time between the pup was put in the cage and the time the vole picked up the pup in its jaws. ‘Groom pup’ was defined as a vole combed the pup’s body surface with its muzzle, accompanied by a rhythmic up-and-down bobbing of the vole’s head and displacement of the pup. ‘Crouch’ was defined as the vole squatted quietly over the pup with no apparent movement. Pup-directed behaviors in optogenetic experiments were scored and analyzed by JWatcher.

### OT treatments

Test virgin voles were acclimatized in their cages for 20 min before being injected intraperitoneally with 0.9% NaCl (1 ml/kg) and pups were introduced 30 min later. The behavioral responses were recorded from the side using a video camera. Thirty mins after the first record, voles were re-injected intraperitoneally OT 1 mg/kg (Leuner, Caponiti, & Gould, 2012) (Bachem, 50-56-6), and pups were introduced 30 min later and behavioral responses were recorded. All behaviors were scored and analyzed using JWatcher.

### Statistics

Parametric tests were used to analyze normally distributed data, and nonparametric tests were used for data that is not normally distributed according to Kolmogorov–Smirnov tests. Independent samples t-tests (two-tailed) were performed to assess number of OT, c-fos and merge rate of OT & c-fos during different pup-directed behaviors (Pup-care vs. Infanticide) and the number of c-fos-IR positive neurons (mCherry vs. CHR2; mCherry vs. eNpHR 3.0). The behavioral changes following optogenetic activation and inhibition (factors: treatment × stimulus) of PVN and mPFC were analyzed by two-way repeated-measures ANOVA. One way ANOVA were used to analyze the AUC changes recorded by the OT sensors in different behaviors. Paired-samples t-test (two-tailed) and Wilcoxin signed ranks test were used to analyze changes in pup-directed behaviors before and after intraperitoneal injection of OT. The Pearson chi-square test was used to compare the difference in the number of infanticide voles between the saline and OT groups. All data were presented as mean ± SEM, statistical analyses of data were performed using MATLAB and SPSS 22.0 software.

## Conflict of interest

The authors declare no conflict of interest.

## Fundings

This research was funded by the STI2030-Majior Projects grant number 2022ZD0205101, the National Natural Science Foundation of China grants numbers 32270510 and 31901082, Natural Science Foundation of Shaanxi Province, China grant number 2020JQ-412, China Postdoctoral Science Foundation grant number 2019M653534 and Fundamental Research Funds for Central University grants numbers GK202301012.

## Notes

### Competing Interest Statement

The authors have declared no competing interest.

### Summary of Updates

Linguistic presentation of the article was revised; effect sizes were reported; gender data were compared; added description of method details; added introduction and discussion; added supplementary data and figures; source data updated

